# Discovery of CRBN-dependent WEE1 Molecular Glue Degraders from a Multicomponent Combinatorial Library

**DOI:** 10.1101/2024.05.04.592550

**Authors:** Hlib Razumkov, Zixuan Jiang, Kheewoong Baek, Inchul You, Qixiang Geng, Katherine A. Donovan, Michelle T. Tang, Rebecca J. Metivier, Nada Mageed, Pooreum Seo, Zhengnian Li, Woong Sub Byun, Stephen M. Hinshaw, Roman C. Sarott, Eric S. Fischer, Nathanael S. Gray

**Author notes:** H.R., Z.J., and K.B. contributed equally to this paper.

## Abstract

Small molecules promoting protein-protein interactions produce a range of therapeutic outcomes. Molecular glue degraders exemplify this concept due to their compact drug-like structures and ability to engage targets without reliance on existing cognate ligands. While Cereblon molecular glue degraders containing glutarimide scaffolds have been approved for treatment of multiple myeloma and acute myeloid leukemia, the design of new therapeutically relevant monovalent degraders remains challenging. We report here an approach to glutarimide-containing molecular glue synthesis using multicomponent reactions as a central modular core-forming step. Screening the resulting library identified HRZ-01 derivatives that target casein kinase 1 alpha (CK1α) and Wee-like protein kinase (WEE1). Further medicinal chemistry efforts led to identification of selective monovalent WEE1 degraders that provide a potential starting point for the eventual development of a selective chemical degrader probe. The structure of the hit WEE1 degrader complex with CRBN-DDB1 and WEE1 provides a model of the protein-protein interface and a rationale for the observed kinase selectivity. Our findings suggest that modular synthetic routes combined with in-depth structural characterization give access to selective molecular glue degraders and expansion of the CRBN-degradable proteome.

## Introduction

Targeted protein degradation is a promising therapeutic modality. Most reported protein degrading molecules affect protein levels through chemically induced proximity between intracellular ubiquitin ligases and a protein of interest, causing polyubiquitination and proteasomal degradation of the latter. Bivalent inducers of proximity recruit their targets and E3 ligases by engaging their respective pockets with high affinity ligands, connected by a linker.^1^ On the contrary, molecular glues stabilize interactions between proteins by modifying the interaction surface without necessarily relying on strong binary affinity.

Bivalent small molecules provide an opportunity for rational degrader design by leveraging potent and selective chemical ligands for targets and the ubiquitination machinery. The resulting compounds can be rationally designed but are limited to proteins for which selective chemical binders can be developed. Furthermore, the large size of these molecules often limits cell permeability and metabolic stability. Conversely, molecular glue degraders can engage conventionally undruggable proteins and generally have more favorable pharmacokinetic profiles due to their smaller size. Several examples of serendipitous discovery of molecular glues have been reported.^3,4^ However, unbiased *de novo* design of molecular glue degraders remains challenging and derivatization of known E3 ligase engaging compounds has proven to be a more effective molecular glue discovery strategy.^5–7^

Cereblon (CRBN), a substrate receptor of the CUL4-RBX1-DDB1-CRBN (CRL4^CRBN^) ubiquitin E3 ligase, is the target of glutarimide-containing chemical ligands. Chemical derivatization of the glutarimide scaffold is an established strategy for expansion of the CRBN molecular glue target space. Thalidomide, pomalidomide, and lenalidomide were the first glutarimide-based degraders approved for the treatment of multiple myeloma (MM) and Erythema Nodosum Leprosum (ENL).^2^ Such compounds bind CRBN and induce ternary complexes with neo-substrate proteins, leading to polyubiquitination and degradation of the latter (Figure 1A). The phthalimide and isoindolinone cores of these compounds occupy the cytosol-exposed tri-tryptophan pocket of CRBN, providing a starting point for derivatization. Previous studies have established chemical substitution as a means to bias CRBN towards a broader range of neo-substrates, which allowed development of more selective molecular glue degraders (Figure S1).

**Figure 1.**
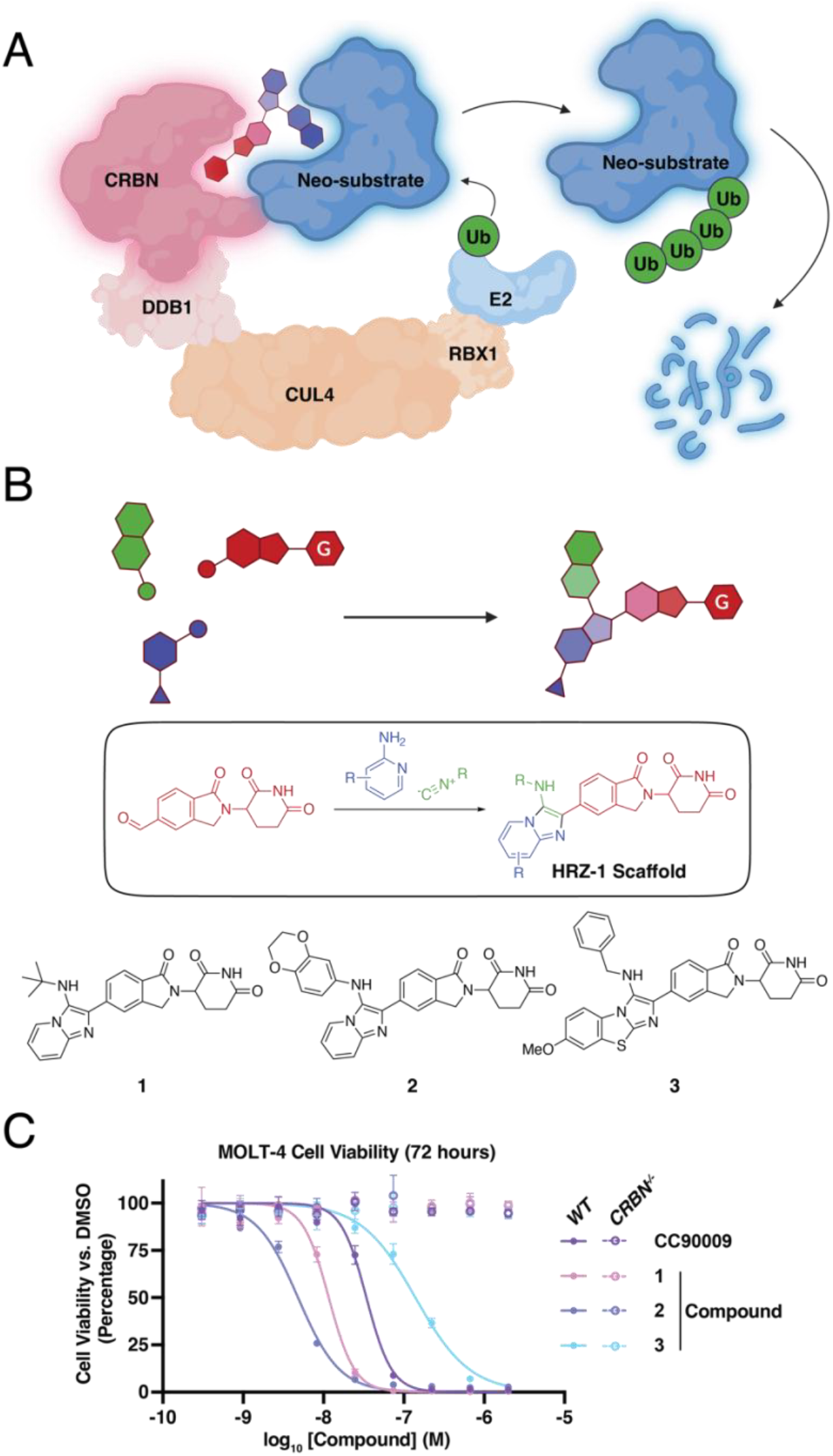
Expanding the chemical space of CRBN molecular glue degraders. (A) Cartoon representation of bivalent (above) and molecular glue (below) degraders mechanisms of action. (B) schematic design of the multicomponent reaction-based library with incorporation of glutarimide moiety (above) and utilization of the GBB reaction for 1-step HRZ-1 scaffold synthesis. (C) Primary MCR library hits of the anti-proliferative phenotypic screening. MOLT-4 cells were incubated with a compound, DMSO as a negative control, or CC-90009 GSPT1 degrader as a positive control at the indicated range of concentrations (n = 3). After 3 days, cell proliferation was evaluated by the Cell-Titer Glo (CTG) assay.

Following the strategy described above, our group developed selective molecular glue degraders through quantitative proteomics profiling of diversity-oriented combinatorial libraries containing a glutarimide pharmacophore. Exploration of the chemical space by sequential substitution of the isoindolinone or another heterocycle attached to glutarimide yielded selective degraders of the phosphodiesterase 6D (PDE6D), CK1α,^8^ Helios zinc finger (IKZF2),^9^ and G1 to S phase transition protein 1, (GSPT1)^10^. Similar approaches have produced clinical stage GSPT1^11,12^ and IKZF1/3 degraders for the treatment of acute myeloid leukemia, multiple myeloma, and non-hodgkin lymphoma [NCT04336982, NCT04756726]. Dual degraders targeting IKZF1/3,^15,16^ CK1α/IKZF2,^17,18^ and GSPT1/2^19,20^ have been reported, and their anti-neoplastic activities have been validated *in vitro* and *in vivo*.

Most glutarimide derivatization strategies rely on stepwise chemical couplings resulting in linear and flexible drug candidates. Linear substitution of the isoindolinone or its isosteres limits chemical space exploration. Additionally, upon identification of a hit degrader derived via this strategy, investigation of the steep structure-activity relationship (SAR) oftentimes relies on laborious stepwise reactions limited to sequential substituent chains generated from a range of available building blocks. Furthermore, the presence of the sensitive glutarimide moiety combined with the stepwise synthetic approach leads to a narrow range of compatible reactions and low total yields. Therefore, improved chemical synthesis strategies for diversifying glutarimide conjugates are highly sought-after.

To circumvent these issues, we devised a synthetic protocol for rapid and efficient access to a new CRBN molecular glue chemical space. We used a Groebke-Blackburn-Bienaymé (GBB)^21^ reaction as the final core-forming step in the synthesis of novel molecular glues. We developed an efficient assay cascade consisting of cell proliferation assays, global proteomics, and subsequent target validation experiments for facile expansion of the CRBN-mediated targeted protein degradation target area as demonstrated by the discovery of a chemical scaffold (HRZ-1) and derivatives that degrade WEE1 and CK1α in a CRBN- and proteasome-dependent manner. Two selective WEE1 degraders were identified. The structure of the ternary complex induced by the most potent hit confirmed binding to the β-hairpin region of the WEE1 kinase domain and helped rationalize the observed selectivity trends. The most promising compounds induced WEE1 degradation and produced anti-proliferative effects in several human cancer cell lines.

## Results

### Multicomponent reaction enables facile synthesis of a diverse CRBN molecular glue library

We used the GBB three-component reaction to synthesize a library of glutarimide derivatives (Figure 1B). The reactants comprised a set of common functionalized glutarimide precursors as the aldehyde components of the GBB reaction. Combining this starting material with commercially available amidines and isocyanides in the presence of a mild to moderately strong Lewis or Brønsted acid resulted in one-pot formation of a wide range of fused heterocyclic cores. By concluding the synthetic route with the GBB reaction, we streamlined the synthesis and obtained over 200 glutarimide derivatives from a handful of glutarimide-containing aldehydes. This approach allows facile synthesis of products with easily modifiable drug-like cores, thus expanding the accessible chemical space for structure-activity relationship (SAR) studies upon hit identification.^22^

### Identification of potent WEE1/CK1α dual degraders

To evaluate over 200 GBB-derived glutarimide derivatives, we established an assay pipeline to identify compounds that induce the degradation of proteins essential for cancer cell viability. Taking advantage of the reported CRBN-dependency of glutarimide derivatives, we carried out 72 hour cell viability experiments in *CRBN* wild-type and knockout MOLT-4 cells (acute lymphoblastic leukemia) to select for compounds that induced potent anti-proliferative effects in a CRBN-dependent manner. Three hits derived from a common aldehyde precursor (HRZ-1 scaffold, Figure 1B, and Table 1) demonstrated potent CRBN-dependent cytotoxicity, reaching half maximal effective concentrations (EC_50_) as low as 5 nM (compound **2**; Figure 1C). Compounds **1** and **3** also had a significant anti-proliferative effect (EC_50_ = 12 and 139 nM, respectively).

**Table 1.**
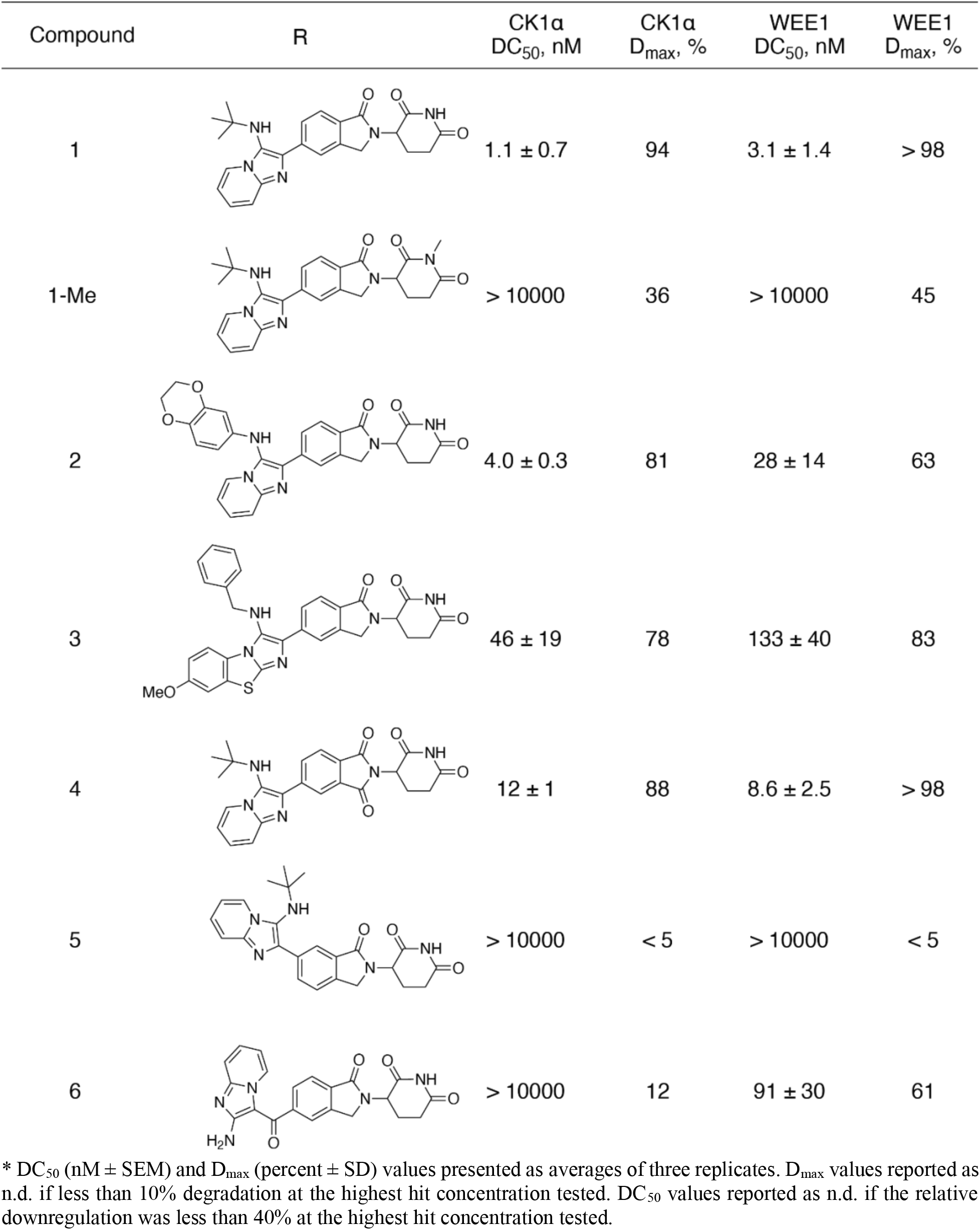
Structure-activity relationship of WEE1/CK1α molecular glue degraders*

| Compound | R | CK1α<br>DC<sub>50</sub>, nM | CK1α<br>D<sub>max</sub>, % | WEE1<br>DC<sub>50</sub>, nM | WEE1<br>D<sub>max</sub>, % |
| --- | --- | --- | --- | --- | --- |
| 1 | | 1.1 ± 0.7 | 94 | 3.1 ± 1.4 | > 98 |
| 1-Me |  | > 10000 | 36 | > 10000 | 45 |
| 2 | | 4.0 ± 0.3 | 81 | 28 ± 14 | 63 |
| 3 | | 46 ± 19 | 78 | 133 ± 40 | 83 |
| 4 | | 12 ± 1 | 88 | 8.6 ± 2.5 | > 98 |
| 5 |  | > 10000 | < 5 | > 10000 | < 5 |
| 6 | | > 10000 | 12 | 91 ± 30 | 61 |

\* DC<sub>50</sub> (nM ± SEM) and D<sub>max</sub> (percent ± SD) values presented as averages of three replicates. D<sub>max</sub> values reported as n.d. if less than 10% degradation at the highest hit concentration tested. DC<sub>50</sub> values reported as n.d. if the relative downregulation was less than 40% at the highest hit concentration tested.

To identify the targets of compound **2**, we used whole-cell proteome profiling of MOLT-4 cells and found that CK1α and WEE1 were the two primary targets after 5 hour treatment in MOLT-4 cells (Figure 2A). Western blotting confirmed CRBN-dependent CK1α and WEE1 degradation in MOLT-4 cells for not only compound **2**, but also compound **1** and **3**. (Figure 2B). Compounds **1** and **3** demonstrated stronger degradation of WEE1 than compound **2**. Pretreatment with MLN4924, an inhibitor of Cullin ligase-dependent ubiquitination, significantly rescued both WEE1 and CK1α degradation induced by compounds **1**-**3** (Figure 2C and S3).

**Figure 2.**
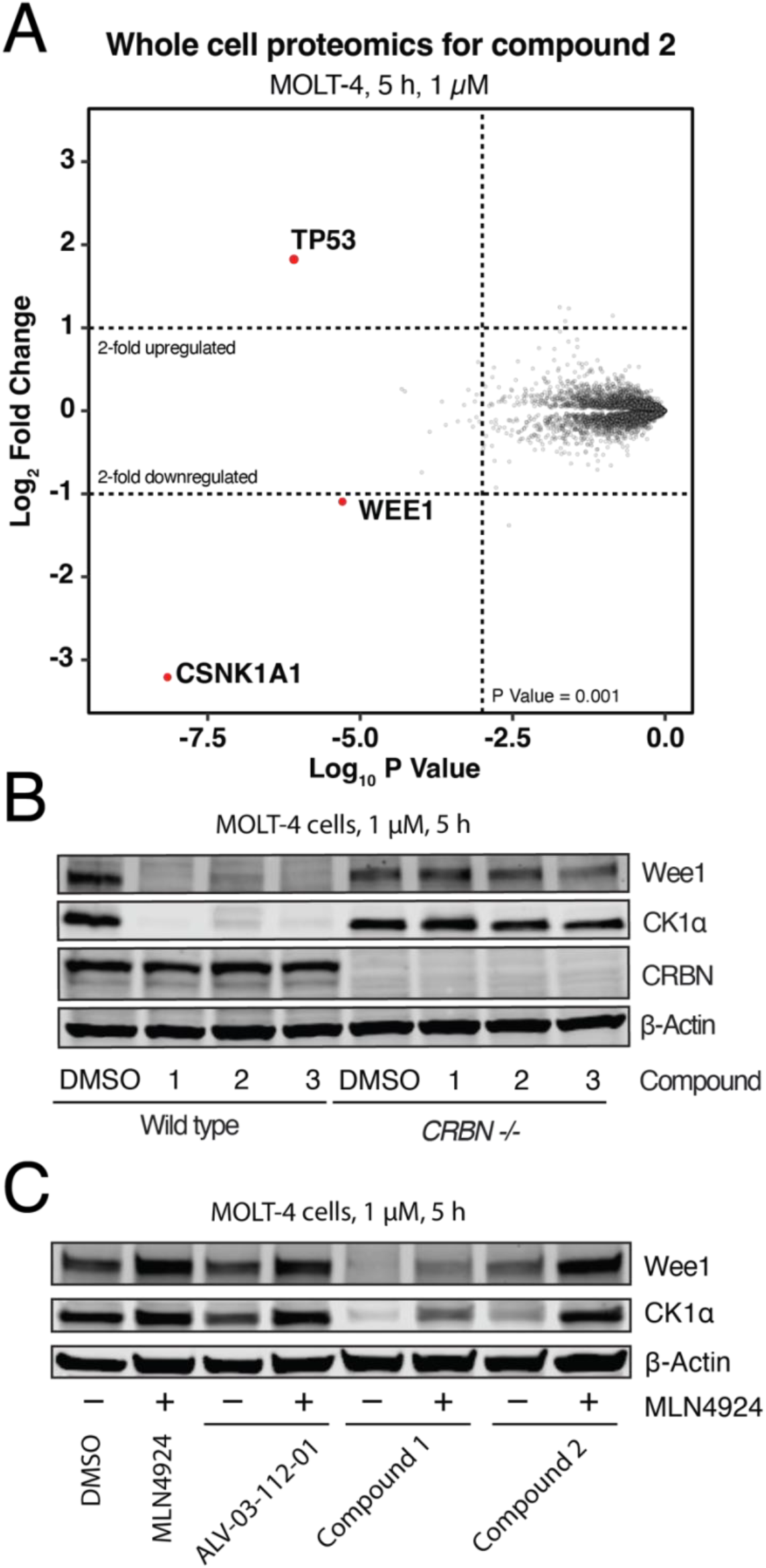
WEE1 and CK1α (also known as CSNK1A1) are degraded by compounds 1-3 in a CRBN E3 ligase-dependent manner. (A) Whole cell proteomics analysis of compound **1**. (B) Western blot of WEE1 and CK1α in wildtype and *CRBN* –/– MOLT-4 cells treated for 5 hours with 1µM compounds **1**-**3**. (C) Western WEE1 and CK1α degradation under in MOLT4 cells pretreated with MLN4924 (1 µM) for 1 hour, followed by treatment with compound **1** and **2** for 5 hours.

To examine their selectivity, we profiled the three hits by Western blot for known CRBN neo-substrates (Helios (IKZF2), Ikaros (IKZF1), and GSPT1, Figure S2). Compound **2** does not induce degradation of any of the tested proteins apart from WEE1 and CK1α at 5 hours. Compound **3** demonstrates modest GSPT1 degradation, which might contribute to its anti-proliferative activity. Compound **1** induces potent degradation of both WEE1 and CK1α and causes incomplete degradation of Helios but not Ikaros. In summary, these results indicate that the identified hits induce WEE1 and CK1α degradation via an established CRBN E3 ligase pathway and have variable activities against known CRBN neo-substrates.

### Structure-activity investigation of HRZ-1 scaffold derivatives neo-substrate selectivity

To explore the structure-activity relationship (SAR) for WEE1 degradation, we used the modular nature of the GBB-based synthetic route to synthesize a range of derivatives. We chose compound **1** as the starting point for the derivatization because it exhibited strong WEE1 degradation while sparing another broadly cytotoxic target, GSPT1 (Figure S2). To facilitate SAR studies, we introduced HiBiT tags^26^ to monitor endogenous WEE1 and CK1α protein levels in Jurkat cells ((T cell leukemia); Figure S4). The modified cell lines were incubated with compounds at a range of concentrations for 5 hours before measurement of target degradation by an endpoint luciferase assay (see Methods).

Screening of the original three hits was in accordance with the western blot results described above. Compound **1** induced WEE1-HiBIT and CK1α-HiBiT degradation at single digit-nanomolar levels, reaching a maximum target fraction degraded (D_max_) of over 98% (Table 1 and Figure S5). No degradation was observed upon treatment with methylated glutarimide **1-Me**, which cannot bind CRBN. In agreement with the western blot results in Figure 2B, compound **2** was a more potent degrader of CK1α than of WEE1. Compound **3** demonstrated less potent half-maximal degradation (DC_50_) values than compound 1 (46 nM vs 1.1 nM for CK1α and 140 vs 3.1 nM for Wee1 respectively).

Compound **1** was selected as the starting point for the SAR due to its promising CRBN-dependent cytotoxicity, potent WEE1 degradation, and absence of GSPT1 degradation (Figure S2, S5). Scaffold hopping from isoindolinone to phthalimide disproportionately weakened CK1α degradation (compound **4**). Switching of the location of the imidazopyridine substitution on the isoindolinone core from C5 to C6 led to a complete loss of CK1α/WEE1-HiBiT degradation (compound **5**). The necessity of the direct C–C bond connection was explored by insertion of a carbonyl linking C5 of the isoindolinone core to C3 of the 2-amino imidazopyridine moiety (compound **6**). Despite a decrease in degradation potency, compound **6** demonstrated pronounced selectivity for WEE1 degradation versus CK1α in the HiBiT assay (Table 1 and Figure S5).

We next explored the amine substituent of compound **1**, which led to the most significant change in selectivity without significant loss of degradation potency (compound **1** vs **2**, Table 1 and Figure S5). Switching the amine to a chloride resulted in compound **7**, which retained strong degradation of both WEE1 and CK1α (compound **7**, Table 2 and Figure S6). Absence of an alkyl substituent (compound **8**) resulted in weaker DC_50_ values for both targets. A phenyl substituent in the same position improved degradation efficiency (compound **9**). The pronounced difference in selectivity between compounds **2** and **9** suggests that the benzodioxane ring is responsible for the observed CK1α preference. A compound bearing a bulky adamantyl substituent (compound **10**, Table 2 and Figure 3A, B) was one of the strongest degraders tested, with both WEE1 and CK1α-HiBiT DC_50_ values reaching sub 2 nM. The degradation of WEE1 by compound **11** was significantly rescued by MLN4924 (Figure S3). Branched hydrophobic substitution (compound **11**) was required to achieve a minor preference for the WEE1-HiBiT degradation.

**Table 2.** SAR profiling of HRZ-1 scaffold aniline substituents *

| Compound | X | Y | R | CK1α<br>DC<sub>50</sub>, nM | CK1α<br>D<sub>max</sub>, % | WEE1<br>DC<sub>50</sub>, nM | WEE1<br>D<sub>max</sub>, % |
| --- | --- | --- | --- | --- | --- | --- | --- |
| 7 | H | CH<sub>2</sub> | Cl | 10.6 ± 0.9 | 84 | 10 ± 8 | 87 |
| 8 | H | CH<sub>2</sub> | NH<sub>2</sub> | 26 ± 3 | 86 | 18 ± 4 | 97 |
| 9 | H | CH<sub>2</sub> |  | 1.8 ± 0.3 | 83 | 1.9 ± 1.1 | 78 |
| 10 | H | CH<sub>2</sub> |  | 1.6 ± 0.5 | 92 | 1.5 ± 0.8 | > 98 |
| 11 | H | CH<sub>2</sub> |  | 9.1 ± 4.3 | 71 | 2.0 ± 0.8 | 98 |
| 12 | H | CO |  | 170 ± 30 | 64 | 22 ± 5 | > 98 |
| 13 | Me | CO |  | > 10000 | < 5 | 140 ± 20 | 85 |

\* DC<sub>50</sub> (nM ± SEM) and D<sub>max</sub> (percent ± SD) values presented as averages of **three replicates**. D<sub>max</sub> values reported as n.d. if less than 10% degradation at the highest hit concentration tested. DC<sub>50</sub> values reported as n.d. if the relative downregulation was less than 40% at the highest hit concentration tested.

**Figure 3.**
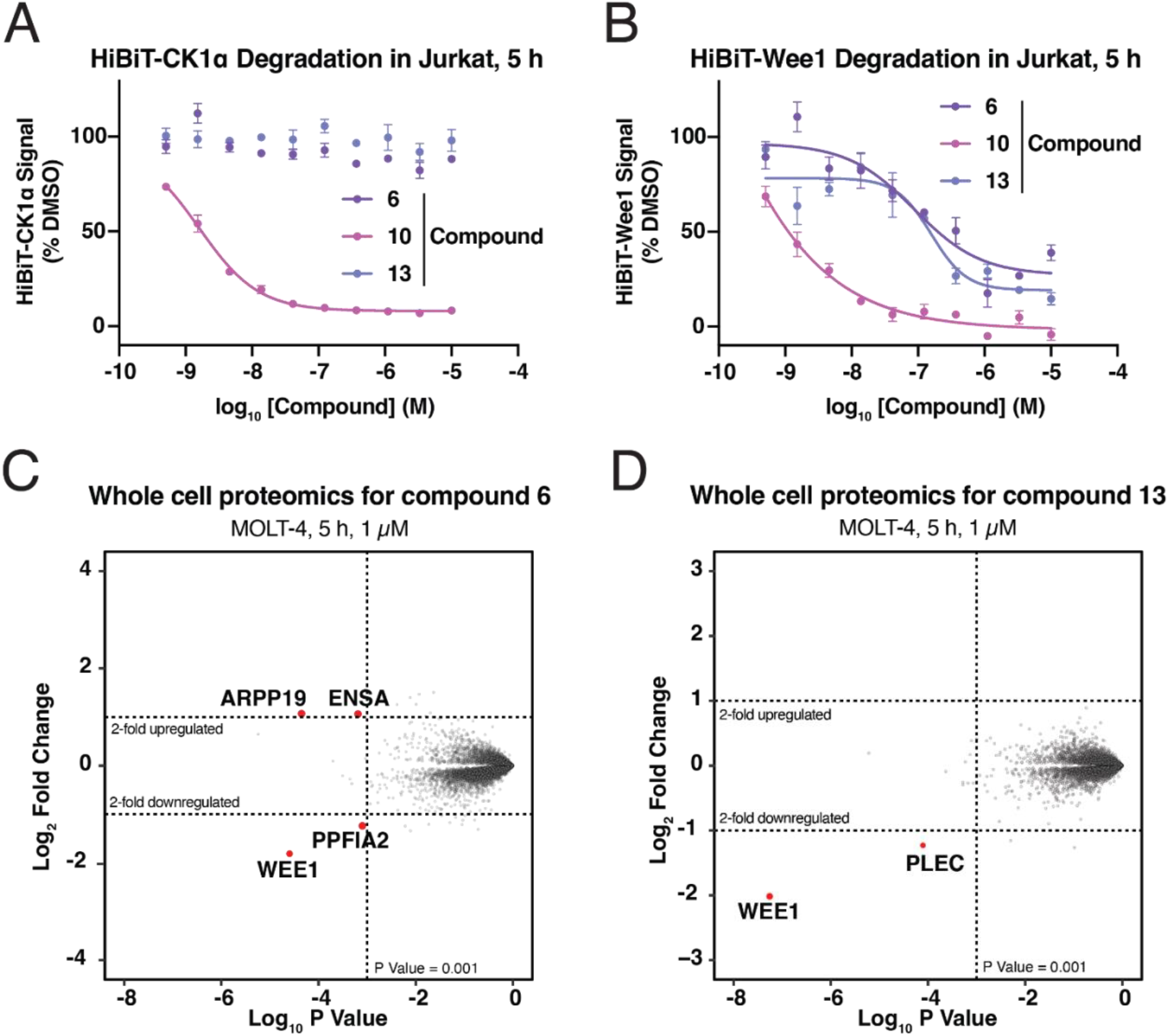
WEE1 degradation selectivity validation. Selective degradation of WEE1 by compounds 6 and 13 were confirmed using HiBiT lytic assay after 5 hour incubation. Results of the assay in Jurkat HiBiT-CK1α (A) and HiBiT-WEE1 (B) cells. Compound **10** was used as a positive control in both experiments. Mean HiBiT signal is presented for three replicates (error bars – ± SD). Compound **6** (C) and **13** (D) were incubated with MOLT-4 cells for 5 hours and the whole cell proteomics analysis was performed.

The SAR indicated that two moieties among the HRZ-1 scaffold derivatives contributed to their preference for WEE1 degradation over CK1α. First, switch to the isoindolinone from phthalimide in compound **4** was more favorable for WEE1 degradation. Second, extension of the hydrophobic amine substituent in compound **11** resulted in a moderate selectivity bias towards WEE1 when compared to its truncated analogue (compound **1**). We combined these moieties in compound **12**, which showed further improvement in WEE1-HiBiT degradation selectivity. We assumed that the amine substituent of compound **12** may be interacting with a lipophilic pocket of WEE1 that is absent or not accessible in CK1α. Introduction of a methyl substituent in a vicinal position to the amine (compound **13**) yielded a selective WEE1-HiBiT degrader (Table 2 and Figure 3A, B), although the selectivity improvement came at the cost of decreased potency. Proteomics experiments confirmed the proteome-wide selectivity of compounds **6** and **13** in MOLT-4 cells after 5 hour treatment at 1 µM (Figure 3C and D).

Overall, the SAR data indicates that linkage of the aromatic systems determines WEE1 selectivity. Additionally, hydrophobic substitutions at the amine affect both potency and selectivity of WEE1- and CK1α-HiBiT degradation. Extended saturated substituents (compounds **11**–**13)** favor WEE1 degradation, while fused ring systems (compound **2**) are biased towards CK1α degradation.

### Structure of the WEE1-compound 10-CRBN complex

We next explored the biochemical mechanism of WEE1 degradation by the most potent degraders (compounds **1** and **10**). To measure compound-induced WEE1-CRBN binding *in vitro*, we used time-resolved fluorescence resonance energy transfer assays (TR-FRET; Figure 4A). The previously reported WEE1 PROTAC, ZNL-02-096^24^,showed a FRET signal peak at ∼10 nM, confirming the sensitivity of the assay. By contrast, even high concentrations of compounds **1** and **10** did not saturate the FRET signal, consistent with a cooperative mode of WEE1-CRBN ternary complex formation. Compound **10** produced a higher FRET signal than compound **1** (4.2 vs 3.2 DMSO-normalized TR-FRET signal ratio). As expected, a weaker WEE1 degrader (compound **2**) did not induce strong FRET signal (< 1.5 fold increase).

**Figure 4.**
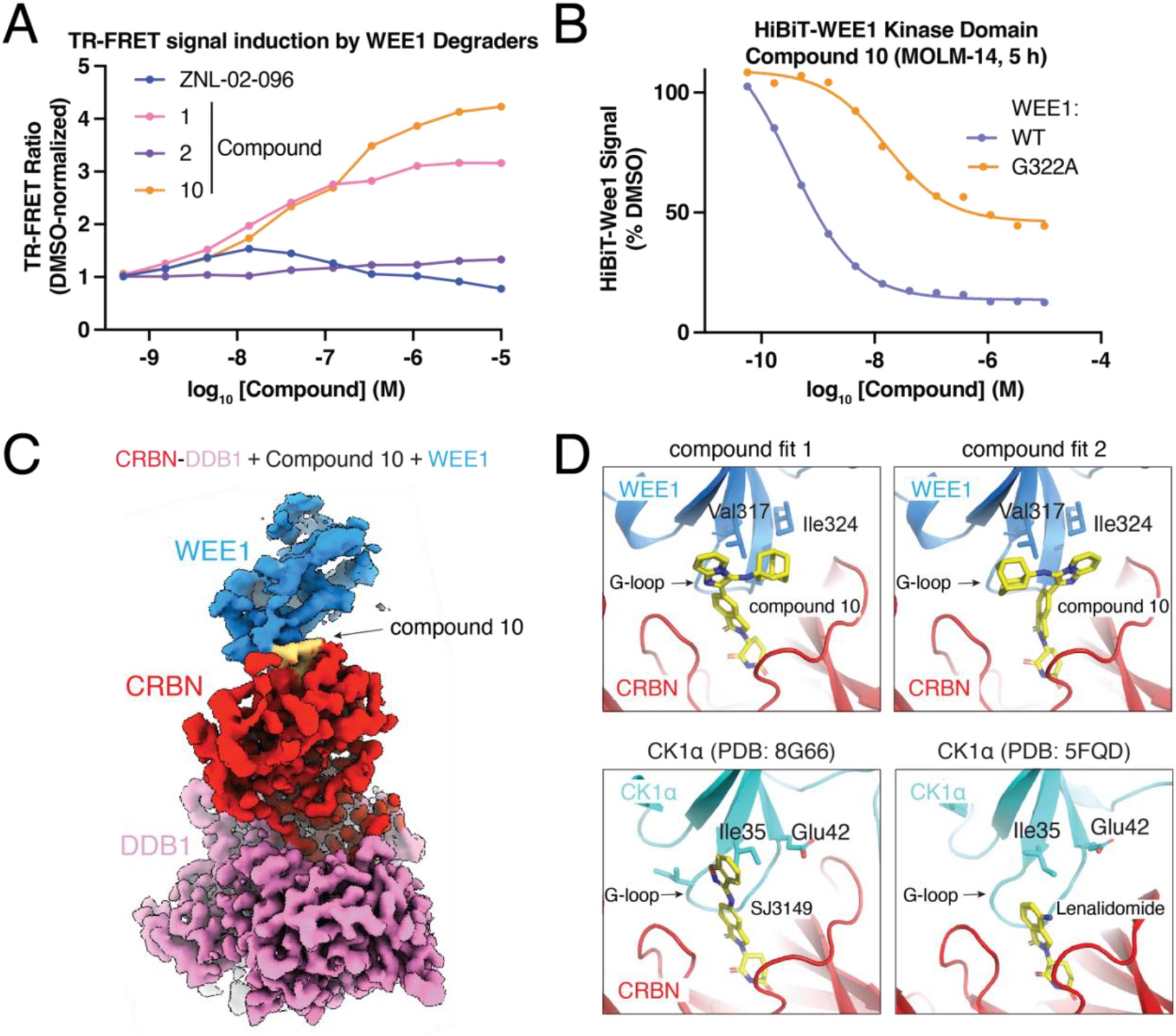
Structural characterization of WEE1 recruitment to CRBN facilitated by compound **10**. (A) Evaluation of TR-FRET signal induced by the hit compounds at a range of concentrations. (B) Mutation of the G-loop glycine leads to rescue of a HiBiT-WEE1 kinase domain degradation by compound 10. (C) Density map of the DDB1-CRBN-compound 10-WEE1 complex visualized by Cryo-EM (left) and magnified WEE1-CRBN interaction region with two possible compound binding modes (right). (D) Comparison of the binding modes of compound 10 and SJ3149 in the CRBN-kinase interaction pocket.

To date, all reported CRBN molecular glues engage neo-substrates via a degron motif composed of a β-hairpin loop containing a glycine residue (G-loop).^27–29^ Except for the invariant presence of a well-positioned glycine, the amino acid sequence of the G-loop is not conserved among known neo-substrates.^30^ The N-lobe of the WEE1 kinase domain contains a β-hairpin moiety centered around glycine 322. To test the hypothesis that this loop is the degron recognized by the HRZ-1 scaffold compounds, we overexpressed WEE1-HiBiT kinase domain (wildtype and G322A) in MOLM-14 cells (Acute Monocytic Leukemia) and measured its abundance after treatment with compound **10** for 5 hours. Compound **10** demonstrated potent degradation of the WT WEE1 kinase domain (DC_50_ = 0.34 nM, D_max_ = 88%), while the G322A mutation showed 50-fold rescue (DC_50_ = 16 nM, D_max_ = 54%) (Figure 4B).

We used cryo-electron microscopy (cryo-EM) to gain structural insights to how compound **10** mediates recruitment of WEE1 to CRBN-DDB1 (Figure 4C and Figure S7). The results show ternary complex formation of WEE1-compound **10**-CRBN-DDB1 complex through the predicted G-loop of WEE1. Rather than a singular stable conformation, multiple WEE1 poses relative to CRBN-DDB1 could be identified, suggesting that despite the high affinity, compound **10** engagement does not rigidify the overall complex (Figure S8A). Furthermore, the heterogeneity of alternating conformations of CRBN, between its open and closed form, precluded us from obtaining a singular stable reconstruction at high resolution (Figure S8B).^31^

Albeit the conformational heterogeneity, the structure is of sufficient quality to enable docking of compound **10** between the WEE1 N-lobe and the CRBN tri-tryptophan pocket in two possible conformations (Figure 4D). The G322-containing G-loop is positioned to contact the compound **10** in a manner analogous to previously reported CRBN molecular glue substrates.^32,33^ Overall and with respect to the compound itself, the structure of the compound **10**-induced ternary complex aligns with the positioning of the DDB1-CRBN-SJ3149-CK1α complex previously reported^34^ (Figure S9). Both possible compound **10** binding modes position the imidazopyridine and adamantyl substituents orthogonally to the benzioxazole moiety of SJ3149 (Figure 4D).

### WEE1/CK1α molecular glue degraders induce target degradation in multiple tumor cell lines

To further explore the versatility of the HRZ-1 scaffold derivatives, compounds **1**-**13** were tested for their abilities to induce WEE1/CK1alpha degradation in multiple tumor cell lines. Selective WEE1 degraders **6** and **13** demonstrated significant and selective degradation of WEE1 over CK1α in Jurkat, MOLM-14, NB-4 (Acute Promyelocytic Leukemia), and LoVo (Colorectal Adenocarcinoma) cell lines (Figure 5A-E). Compound **13** partially degraded CK1α in SU-DHL-5 cells (Diffuse Large B Cell Lymphoma), while compound **6** maintained WEE1 selectivity in these cells (Figure 5B). Compound **10** was used as a reference for potent degradation of both kinases.

**Figure 5.**
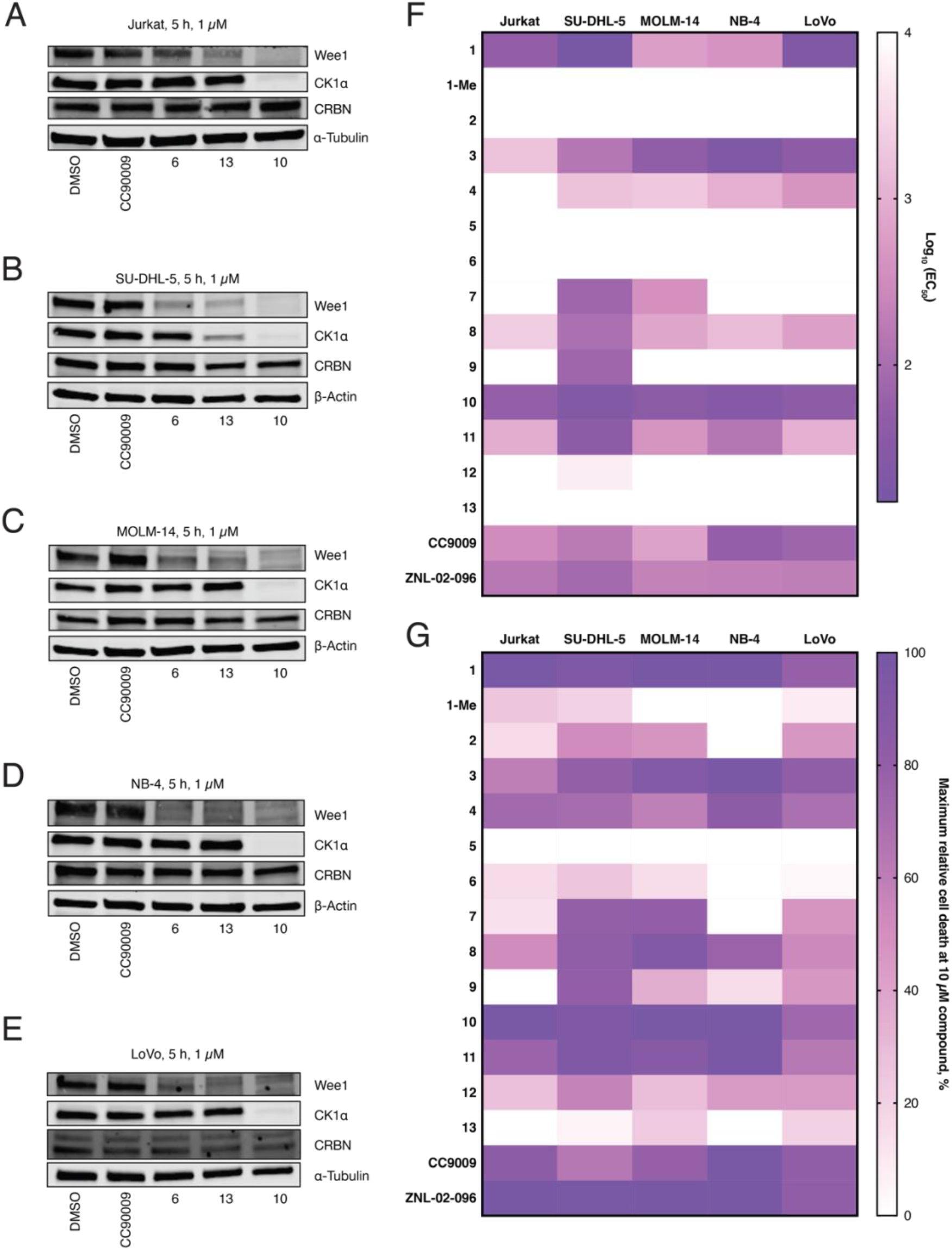
Characterization of the HRZ-1 scaffold derivatives CK1α/WEE1 degradation and anti-proliferative effect in a range of cancer cell lines. (A-E) Western blots showing consistent and preferential degradation of WEE1 by selective degraders **6** and **13**. CC90009 and Compound **10** were used as negative and positive controls, respectively. (F, G) Heat maps demonstrating effect of compounds **1**-**14** on the cancer cells viability using EC_50_ (compound concentration leading to half-maximal cell death) and the maximum relative cell death achieved at 10 µM treatment.

The anti-proliferative effects of compounds **1**-**13** were measured in SU-DHL-5, MOLM-14, NB-4, and LoVo cells. EC_50_ concentrations and the maximum cell death at 10 µM compound relative to the DMSO control were determined (Figure 5F and G). Compounds **1-Me** and **5** were inactive in all tested cell lines, demonstrating that glutarimide-CRBN engagement and a fitting exit vector from the isoindolinone scaffold are required for activity. Compounds **1** and **10** were the most potent antiproliferative compounds, outperforming the GSPT1 degrader CC90009 and ZNL-02-096. The selective compounds induced mild cytotoxic effects in several tested cell lines but did not decrease relative proliferation below 50%. One possible explanation for these cell cytotoxicity results is chemical instability of the glutarimide core, a possibility we are currently investigating.

## Discussion

Here, we report synthetic strategy for facile generation of a chemically diverse library of glutarimide derivatives. GBB reactions yielded drug-like molecular glue candidates from a handful of glutarimide-containing aldehyde intermediates and a range of commercial amidines and isocyanides. We identified CK1α and WEE1 as the primary targets of the GBB-derived HRZ-1 scaffold hits using a phenotypic assay pipeline paired with label-free quantitative proteomics analysis. Both targets are degraded in a CRBN E3 ubiquitination machinery-dependent manner. This approach stands out due to a large focus on increased hit diversification feasibility, while following established post-hit identification mechanism validation steps seen in other CRBN molecular glue discovery studies.^8–10,12,18^ Utilizing multicomponent reaction gives access to pharmacophores that can be diversified with minimal synthetic efforts down the degrader hit-to-lead optimization path.

The SAR expansion of the original hits provided structural insights into WEE1 vs CK1α selectivity trends. The most potent dual degrader, compound **10**, showed sub-nanomolar half-maximal WEE1 degradation. The WEE1 degradation selectivity of compounds **6** and **13** was confirmed by proteome-wide quantitative analysis. These selective hits provide a basis for the future development of potent and selective WEE1 degraders, despite their limited potency and lack of anti-proliferative effects in cancer cell lines.

Biochemical experiments and a cryo-EM structure confirmed the molecular glue-like mechanism of action of HRZ-1-related compounds. G322A mutant of WEE1-HiBiT displayed significantly decreased degradation by compound **10**, suggesting that the G322 is indeed part of the engaged degron, as the previously reported G-loop mutants have led to complete loss of the degrader activity^17,19^. However, compound **10** still leads to mild downregulation of this G-loop mutant protein. One explanation for the observed partial tolerance to alanine could be a more extensive protein-protein interface or structural flexibility tolerating the bulky sidechain of alanine over glycine. The structure of WEE1-compound **10**-CRBN-DDB1 complex supplemented with site-specific glycine mutagenesis validated the G-loop degron in the WEE1 kinase domain and visualized flexible engagement of the kinase to the CRBN tri-tryptophan pocket. The degrader-induced complex structure aligns well with the previously reported CK1α-SJ3149-CRBN-DDB1 structure.^34^ Based on the compound **10** binding mode, the selectivity of compound **13** can be rationalized by broader engagement of the hydrophobic Val317 and Ile324 amino acids in the WEE1 G-loop (Figure 4D), while the corresponding residues in the CK1α ternary complex appear to be more hydrophilic. These steric and electrostatic differences in the respective kinase surfaces help rationalize the observed differences in degraders selectivity and provide a basis for the rational design of more potent and selective WEE1 degraders.

The CK1α/WEE1-targeting HRZ-1 scaffold derivatives showed strong anti-proliferative effects in both hematological and solid tumor cell lines. WEE1 is an essential cell cycle kinase that phosphorylates and inhibits CDK1/Cyclin B in response to DNA damage.^25^ Inhibition of WEE1 activity in malignancies with high genomic instability and dysregulated G1-S checkpoint signaling sensitizes them to DNA-damaging agents.^35^,^25^ The WEE1 inhibitor AZ1775 has shown efficacy in clinical trials, but significant adverse effects have slowed its development.^36^ Our group^24^ and others^37^ have reported proteolysis targeting chimera (PROTAC) WEE1 degraders based on this inhibitor. WEE1 glue compounds, which are expected to have more drug-like properties than previously reported small molecules, have not been described in the peer-reviewed literature and therefore represent a distinct therapeutic opportunity. Compounds **1** and **10** demonstrated comparable or stronger cytotoxic effects compared with the published WEE1 PROTAC ZNL-02-096 and the GSPT1 degrader CC90009. Some of the WEE1/CK1α derivatives with similar degradation potencies have demonstrated comparable effects on cellular viability (compounds **8** and **11**), while others did not induce potent cell death (compounds **7** and **9**). Additionally, the selective WEE1 degraders did not induce strong cell killing. Further investigation will be needed in order to understand the different anti-proliferative effects of reported HRZ-1 derivatives, regardless of WEE1 and CK1α degradation observed. This could be indicative of the broader range of neo-substrates degraded by the cytotoxic compounds. Alternatively, partial degradation of WEE1 and/or CK1α may lead to growth arrest and senescence rather than apoptosis.

## Conclusions

In summary, we have developed an efficient synthetic sequence and assays to develop novel molecular glue degraders of CRBN neo-substrates. Our studies culminated in the discovery of potent and selective degraders of WEE1, an important cell-division kinase that is deregulated in many cancers. In-depth proteomic experiments and cryoEM structures rationalize the observed SAR and serve as blueprint for further improved candidate molecules.

## Supporting information

Cellular and Molecular Biology Methods

Synthetic Methods

## Acknowledgements

We thank the Harvard Cryo-EM Center for Structural Biology, and M. Hunkeler for helpful discussions. This work has been supported by the National Institutes of Health (NIH) grants R01CA262188 and R01CA2144608 (to E.S.F.), R01CA218278 (to N.S.G. and E.S.F.), NIH High End Instrumentation grant 1S10OD028697-01 (to E.S.F.), and departmental funds from Stanford Chemical and Systems Biology and Stanford Cancer Institute (to N.S.G.). K.B. acknowledges Damon Runyon Fellow supported by the Damon Runyon Cancer Research Foundation (DRG-2514-24). R.C.S. acknowledges the Swiss National Science Foundation for a postdoctoral fellowship (SNF Mobility grant P500PN_206898). M.T.T. acknowledges ChEM-H Chemistry/Biology Interface Training Program T32 (grant 5T32GM139791-02).

## Notes

The authors declare the following competing financial interest(s): E.S.F. is a founder, scientific advisory board (SAB) member, and equity holder of Civetta Therapeutics, Proximity Therapeutics, and Neomorph, Inc. (also board of directors). He is an equity holder and SAB member for Avilar Therapeutics, Photys Therapeutics, and Ajax Therapeutics and an equity holder in Lighthorse Therapeutics. E.S.F. is a consultant to Novartis, EcoR1 capital, Odyssey and Deerfield. The Fischer lab receives or has received research funding from Deerfield, Novartis, Ajax, Interline, Bayer and Astellas. N.S.G. is a founder, science advisory board member (SAB) and equity holder in Syros, C4, Allorion, Lighthorse, Voronoi, Inception, Matchpoint, Cobro Ventures, GSK, Larkspur (board member), Shenandoah (board member), and Soltego (board member). The Gray lab receives or has received research funding from Novartis, Takeda, Astellas, Taiho, Jansen, Kinogen, Arbella, Deerfield, Springworks, Interline and Sanofi. M.S. has received research funding from Calico Life Sciences LLC. K.A.D. receives or has received consulting fees from Kronos Bio and Neomoprh Inc.

