## Supplementary Figures and Tables for "Discovery of CRBN-dependent WEE1 Molecular Glue Degraders from a Multicomponent Combinatorial Library"

**Affiliations:**
[1]Department of Chemistry, Stanford School of Humanities and Sciences, Stanford University, CA, 94305, USA
[2]Department of Chemical and Systems Biology, ChEM-H, Stanford School of Medicine, Stanford University, Stanford, CA, 94305, USA
[3]Department of Cancer Biology, Dana-Farber Cancer Institute, Boston, MA, 02215, USA
[4]Department of Biological Chemistry and Molecular Pharmacology, Harvard Medical School, Boston, MA, 02115, USA
[5]Stanford Cancer Institute, Stanford School of Humanities and Sciences, Stanford University, CA, 94305, USA

### Supplementary Figures

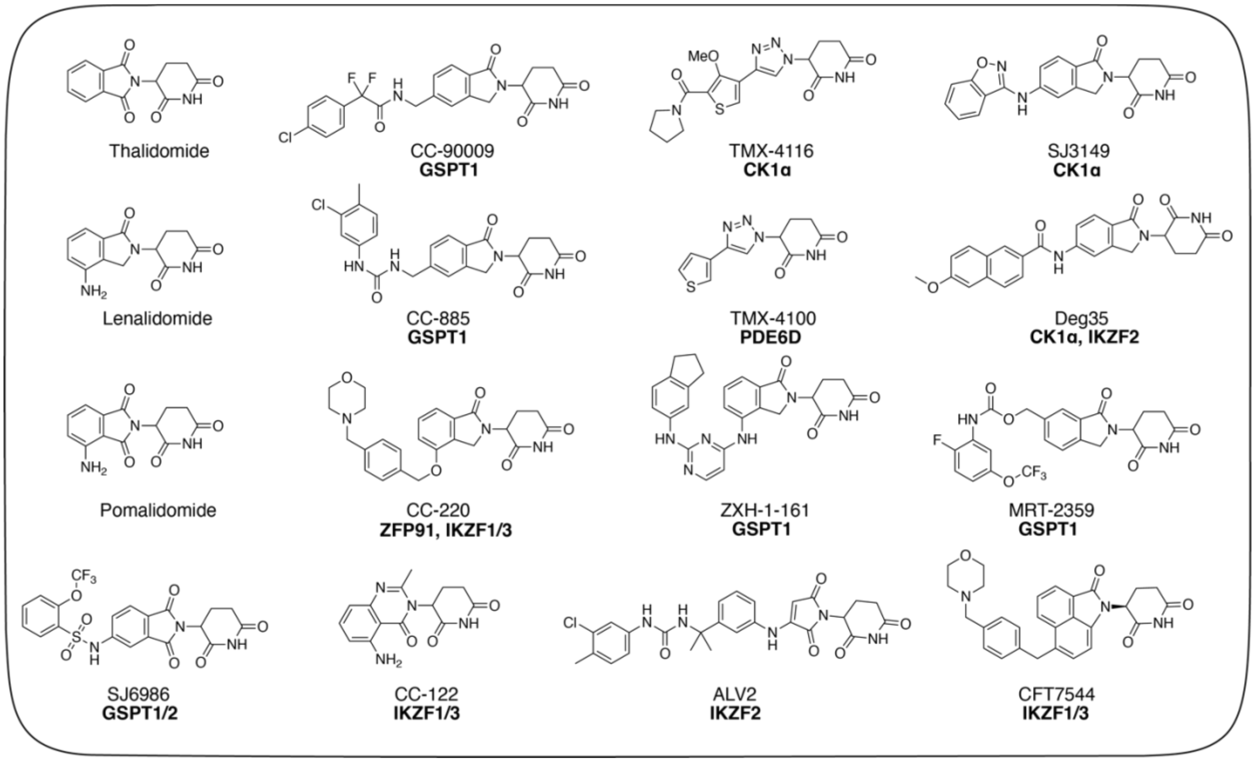


. **Figure S1.** Previously reported CRBN molecular glue degraders and their primary targets.

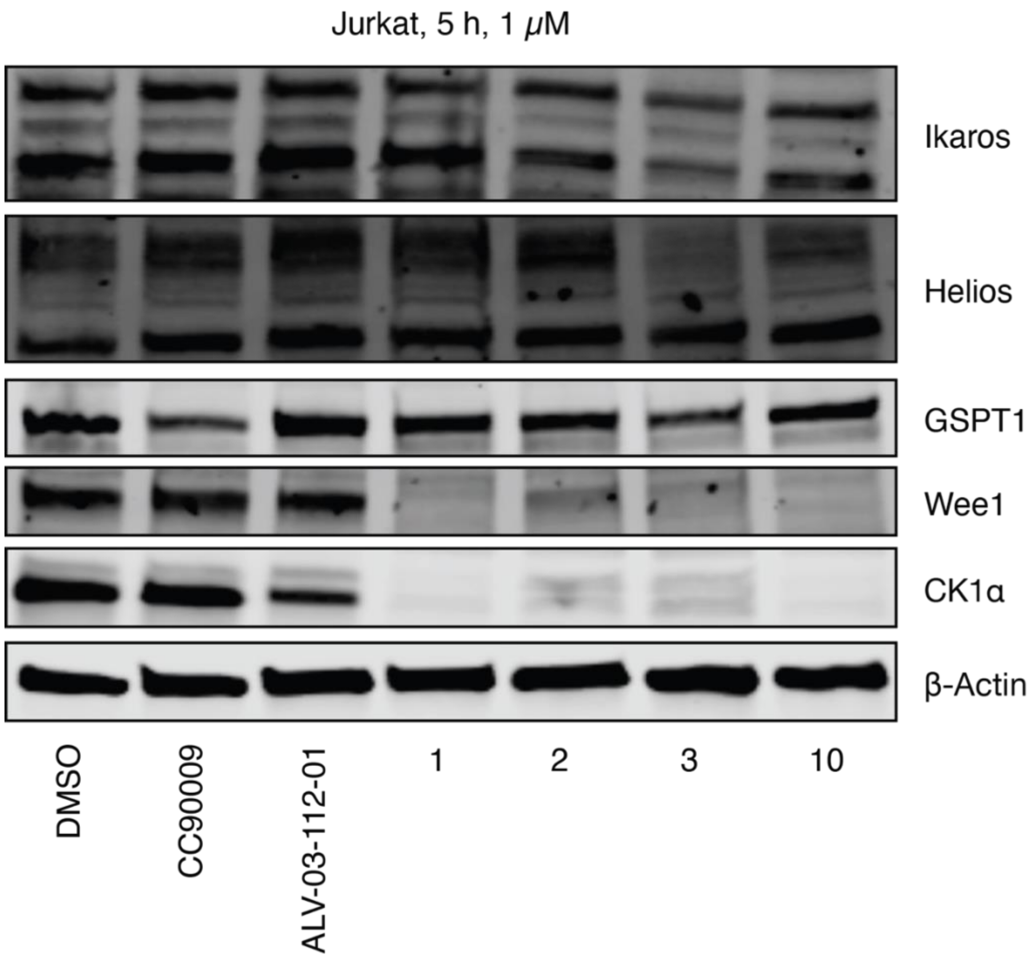


**Figure S2.** Western blot showing degradation effects of compounds **1**-**3** and **10** on the known CRBN neo-substrates. Jurkat cells were incubated with 1 μM of HRZ-1 scaffold hits, Helios degrader ALV-2, or GSPT1 degrader CC-90009 for 5 hours.

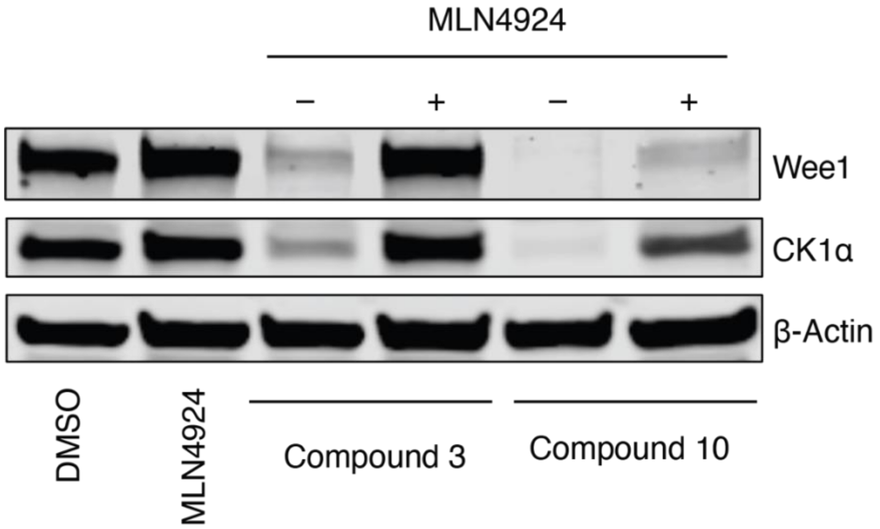


**Figure S3.** Western blot showing rescue of WEE1 and CK1α degradation by compounds **3** and **10** under 1 hour 1 μM pretreatment with MLN4924 that inhibits Cullin-dependent ubiquitination.

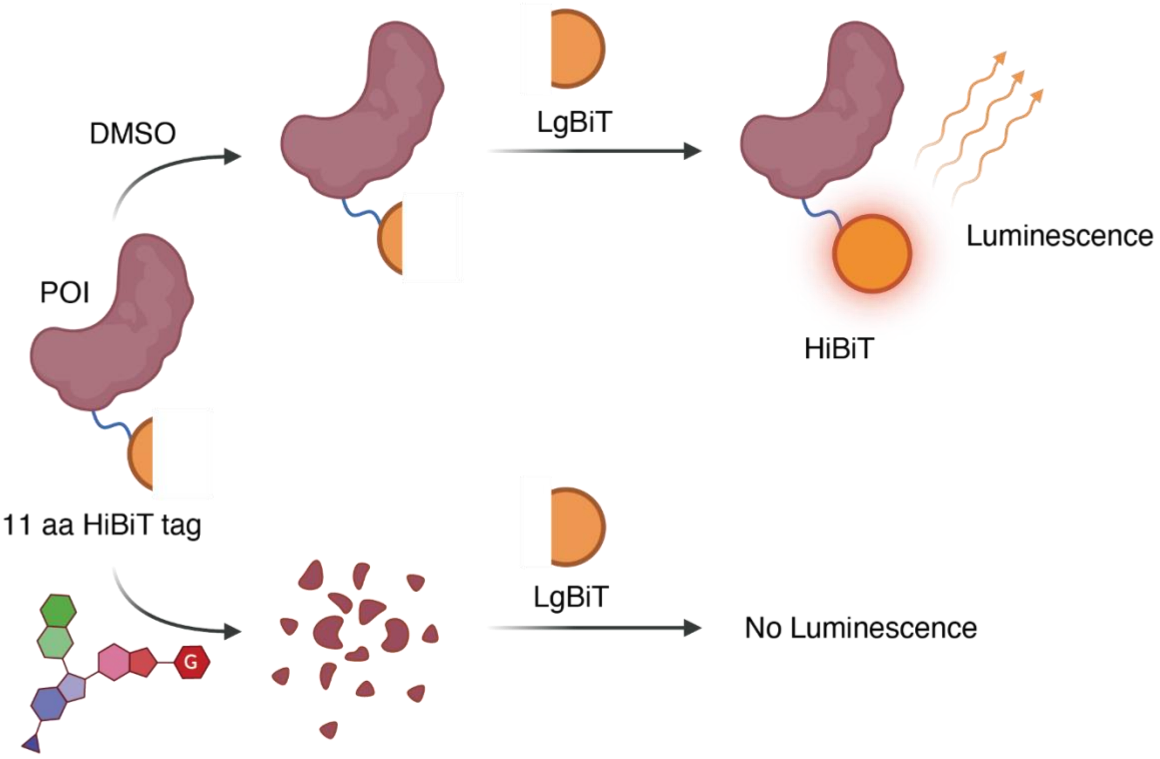


**Figure S4.** Cartoon depiction of the hit modification process. An N-terminal HiBiT tag was knocked into endogenous proteomics hits: WEE1 and CK1α. The resulting HiBiT Jurkat cell lines were used to establish selectivity and potency profiles of the HRZ-1 scaffold derivatives. The obtained selective WEE1 degraders were then tested in a proteomics experiment profiling to establish their proteome-wide selectivity profile.

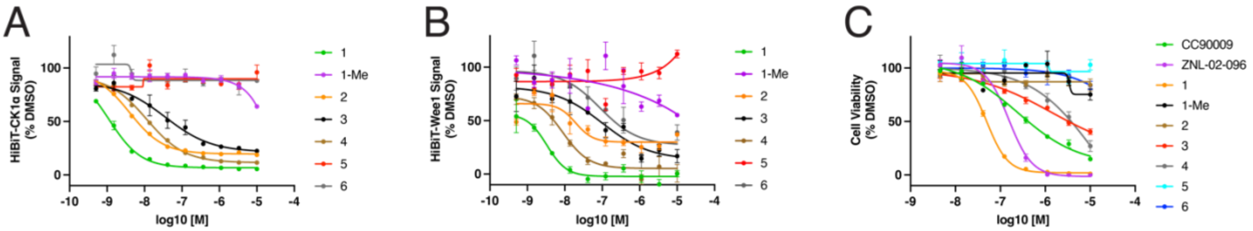


**Figure S5.** Expanded degradation and anti-proliferative activity data for compounds **1**-**6** in Table 1. (A, B) HiBiT-CK1α/WEE1 degradation at a range of concentrations. (C) Anti-proliferative effect of compounds **1**-**6** at a range of concentrations.

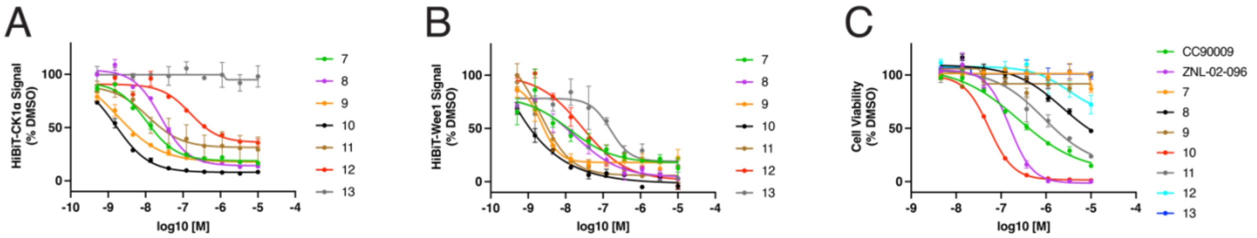


**Figure S6.** Expanded degradation and anti-proliferative activity data for compounds **1**-**6** in Table 2. (A, B) HiBiT-CK1α/WEE1 degradation at a range of concentrations. (C) Anti-proliferative effect of compounds **7**-**13** at a range of concentrations.

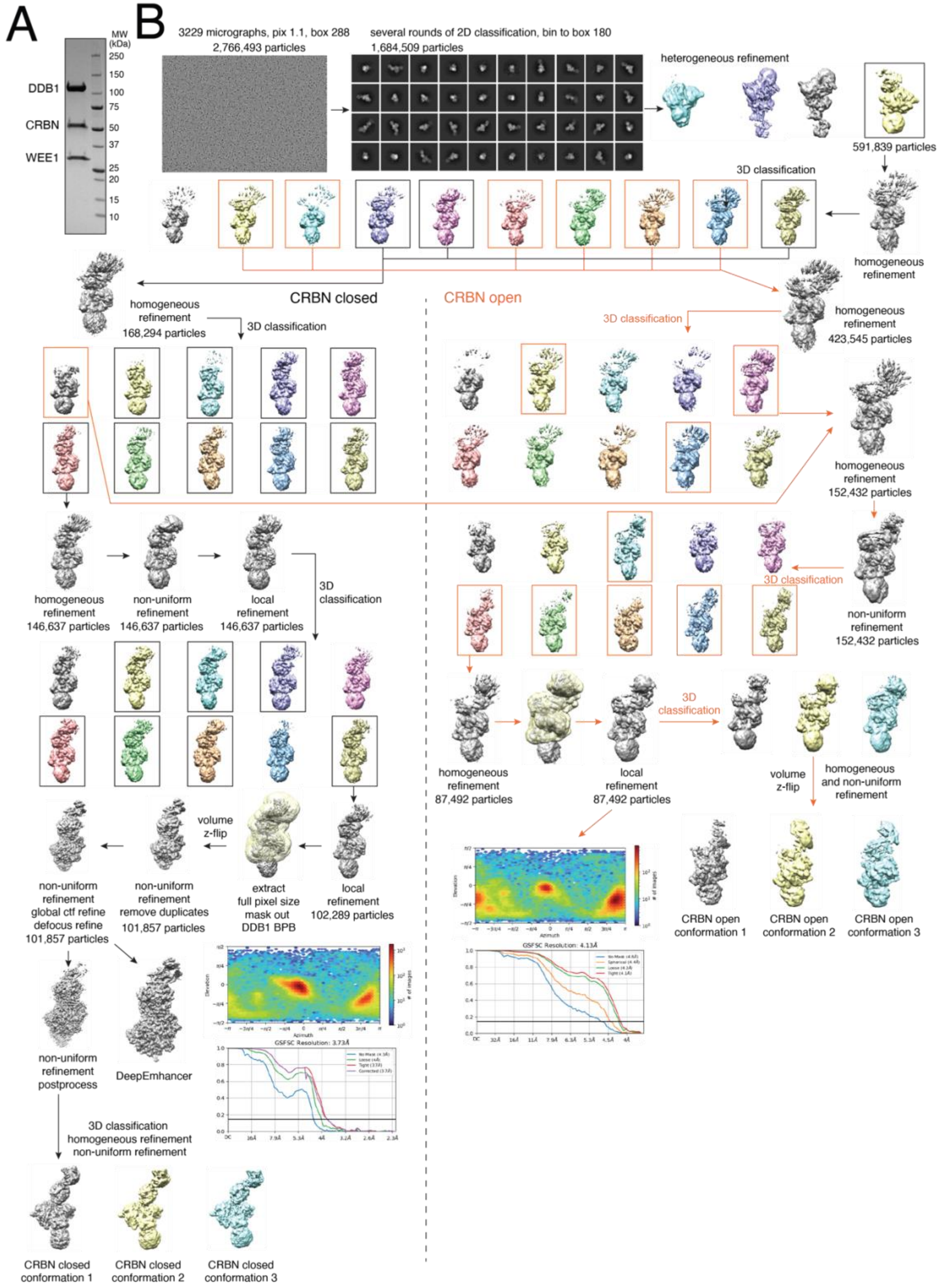


**Figure S7.** Cryo-EM sample preparation and data processing. (A) Coomassie stain of SDS-PAGE showing WEE1-compound 10-CRBN-DDB1 complex. (B) Cryo-EM processing schematic

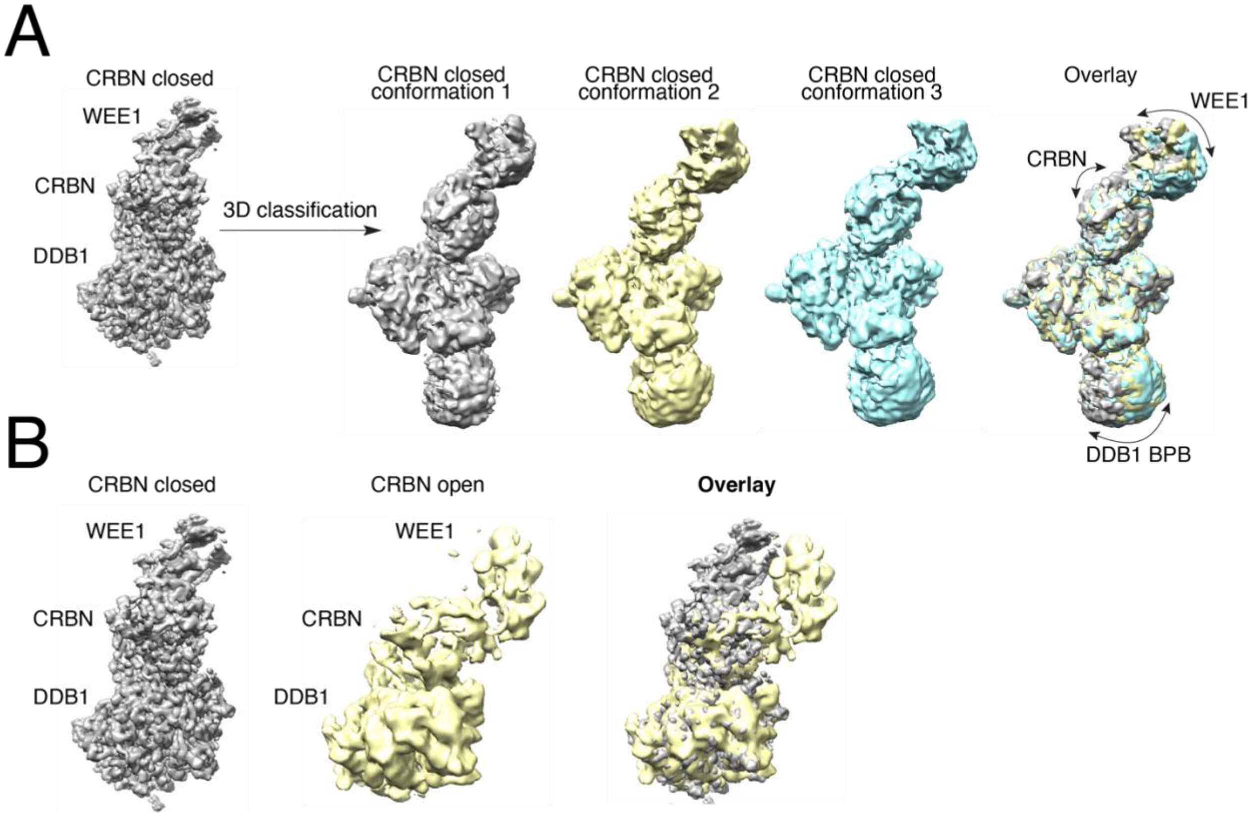


**Figure S8.** Compound 10 brings CRBN and WEE1 in proximity without excessively rigidifying the ternary complex. Density maps of the ternary complex showing (A) high flexibility with the CRBN in closed confirmation and (B) recruitment of WEE1 by compound 10 to both closed and open CRBN confirmations.

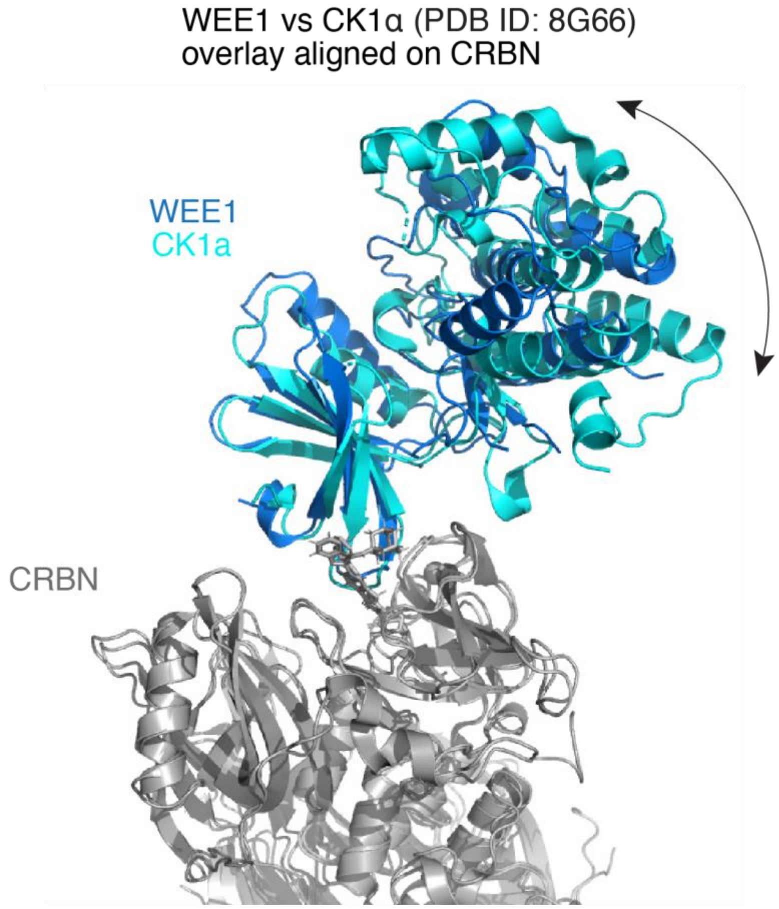


**Figure 9.** Overlay of the protein complex in Figure 4C with the previously reported DDB1-CRBN-SJ3149-CK1α aligned on CRBN

**Table S1.** $EC_{50}$ values determined from the cellular viability experiments in Figure S5, S6, and S10.

| **Compound** | **Jurkat** | **SUDHL5** | **Molm14** | **NB-4** | **LoVo** |
|---|---|---|---|---|---|
| **1** | 49 | 15 | 570 | 400 | 20 |
| **1-Me** | > 10000 | > 10000 | > 10000 | > 10000 | > 10000 |
| **2** | > 10000 | > 10000 | > 10000 | > 10000 | > 10000 |
| **3** | 1700 | 140 | 46 | 21 | 40 |
| **4** | > 10000 | 1600 | 2000 | 1000 | 420 |
| **5** | > 10000 | > 10000 | > 10000 | > 10000 | > 10000 |
| **6** | > 10000 | > 10000 | > 10000 | > 10000 | > 10000 |
| **7** | > 10000 | 72 | 370 | > 10000 | > 10000 |
| **8** | 2300 | 104 | 730 | 1300 | 600 |
| **9** | > 10000 | 72 | > 10000 | > 10000 | > 10000 |
| **10** | 54 | 23 | 35 | 26 | 40 |
| **11** | 900 | 41 | 420 | 140 | 1000 |
| **12** | > 10000 | 5800 | > 10000 | > 10000 | > 10000 |
| **13** | > 10000 | > 10000 | > 10000 | > 10000 | > 10000 |
| **CC9009** | 300 | 170 | 660 | 54 | 71 |
| **ZNL-02-096** | 150 | 84 | 220 | 210 | 190 |

**Table S2.** Maximum antiproliferative effect at 10 µM values determined from the CTG cellular viability experiments in Figure S5, S6, and S10.

| **Compound** | **Jurkat** | **SUDHL5** | **Molm14** | **NB-4** | **LoVo** |
|---|---|---|---|---|---|
| **1** | 99 | 99 | 100 | 100 | 79 |
| **1-Me** | 25 | 22 | -1 | -24 | 9 |
| **2** | 16 | 77 | 47 | 0 | 46 |
| **3** | 60 | 76 | 92 | 98 | 82 |
| **4** | 73 | 97 | 59 | 85 | 69 |
| **5** | -3 | 5 | -5 | -7 | -10 |
| **6** | 16 | 48 | 15 | -8 | 4 |
| **7** | 13 | 98 | 81 | -19 | 47 |
| **8** | 52 | 89 | 93 | 77 | 54 |
| **9** | -1 | 88 | 35 | 15 | 46 |
| **10** | 99 | 98 | 99 | 99 | 74 |
| **11** | 76 | 98 | 89 | 98 | 63 |
| **12** | 28 | 85 | 29 | 45 | 45 |
| **13** | 0 | 83 | 23 | -6 | 20 |
| **CC9009** | 85 | 100 | 79 | 99 | 82 |
| **ZNL-02-096** | 100 | 84 | 100 | 100 | 83 |
