## Supplementary material for "Discovery of CRBN-dependent WEE1 Molecular Glue Degraders from a Multicomponent Combinatorial Library": Cellular and Molecular Biology Methods

**Cellular and molecular biology methods for:**

*Cell line growth and maintenance*

Jurkat, MOLT-4, SU-DHL-5, Molm-14, NB-4 were maintained in RPMI-1640 (Gibco) supplemented with 10% heat-inactivated fetal bovine serum (FBS, Gibco) and 100 U/mL penicillin-streptomycin (Gibco). LoVo were cultured in F-12K (ATCC) supplemented with 10% heat-inactivated FBS (Gibco) and 100 U/mL penicillin-streptomycin (Gibco). All cell lines were cultured at 37 ℃ in a humidified incubator in the presence of 5% $CO_2$. Mycoplasma testing was performed on a monthly basis using the MycoAlert mycoplasma detection kit (Lonza, Basel, Switzerland) and all lines were negative.

*Immunoblotting*

Whole cells lysates for immunoblotting were prepared by pelleting cells from each cell line at 4 ℃ (500 g) for 5 minutes. The resulting cell pellets were washed once with ice-cold 1x PBS and then resuspended in RIPA Lysis and Extraction Buffer (Thermo Fisher Scientific) supplemented with protease and phosphatase cocktails (Roche). Lysates were clarified at 20,000 g for 15 minutes at 4 ℃. Protein concentrations were determined by BCA protein assay (Pierce). Whole cell lysates were loaded into 4-20% precast polyacrylamide gels (Bio-Rad) and separated by electrophoresis. The gels were transferred to a nitrocellulose membrane (Bio-Rad) and blocked for 1 hour at room temperature in Intercept (TBS) Blocking Buffer (LI-COR). Membranes were probed using antibodies raised against Wee1 (Cell Signaling Technology, #13084S), CK1α (Abcam, #ab108296), GSPT1 (Abcam, #ab49878), Ikaros (Cell Signaling Technology, #14859S), Helios (Cell Signaling Technology, #42427S), CRBN (Novus Biologicals, #NBP1-91810), α-Tubulin (Cell Signaling Technology, #3873S), or β-Actin (Cell Signaling Technology, #3700) at 4 ℃ overnight. Membranes were incubated with the IRDye800-labeled goat anti-rabbit IgG or IRDye680-labeled goat anti-mouse IgG (LI-COR) secondary antibodies at room temperature for 1 hour, and detected using an Odyssey CLx system.

*Generation of Wee1/CK1α HiBiT Jurkat cells*

Introduction of a HiBiT coding sequence into the endogenous *Wee1/CK1α* locus in Jurkat cells was done via CRISPR-Cas9 genome editing. ALT-R CRISPR RNA (crRNA) and *trans*-activating CRISPR RNA (tracrRNA) (Integrated DNA Technologies, IDT) were resuspended in Nuclease-Free Duplex Buffer (IDT) at a concentration of 200 μM each. Equal volumes of crRNA and tracrRNA were mixed (final concentration of 100 μM each) and heated for 5 minutes at 95℃. After heating, the complex was gradually cooled to room temperature. The oligo complex was then incubated at room temperature for 20 minutes with the Cas9 Nuclease V3 (IDT) to form the ribonucleoprotein (RNP) complex. The double stranded DNA HDR template (HiBiT-Wee1 with extensions/HiBiT-CK1α with extensions), the RNP complex, and an electroporation enhancer (IDT) were then electroporated into Jurkat cells using an Amaxa 4Dnucleofector (Lonza). Electroporated cells were transferred to medium with HRD enhancer (IDT). Single cells were subsequently isolated via fluorescence activated cell sorting, and HiBiT expression from individual clones was detected with the Nano-Glo HiBiT Lytic Detection System (Promega).

Wee1:
>crRNA sequence
GAGTAGCTCAGTATATAGTA
>HDR donor sequence (+)
GACTTATTGGAAAGAAAATGAACCGCTCTGTCAGCCTTACTATATACGGGAGTTCTG
GCGTGAGCGGCTGGCGGCTGTTCAAGAAGATTAGCTGAGCTACTCCTTTCCCACCTC
CCCCTGAACACTGTGACA
>HDR donor sequence (-)
TGTCACAGTGTTCAGGGGGAGGTGGGAAAGGAGTAGCTCAGCTAATCTTCTTGAAC
AGCCGCCAGCCGCTCACGCCAGAACTCCCGTATATAGTAAGGCTGACAGAGCGGTT
CATTTTCTTTCCAATAAGTC
CK1α:
>crRNA sequence
TCAATTCATGCTTAGAAACC
>HDR donor sequence (+)
ATCATCTGCTCTGCTTCTTCTGTTCCTCAATTCATGCTTAGCTAATCTTCTTGAACAGC
CGCCAGCCGCTCACGCCAGAACTCCCGAAACCTGGTAAAAAATCAAAACAAAAAAA
AAGTTAAAATTCC
>HDR donor sequence (-)
GGAATTTTAACTTTTTTTTTGTTTTGATTTTTTACCAGGTTTCGGGAGTTCTGGCGTGA
GCGGCTGGCGGCTGTTCAAGAAGATTAGCTAAGCATGAATTGAGGAACAGAAGAAG
CAGAGCAGATGAT

*Generation of WEE1 kinase domain-HiBiT wildtype and G322A overexpressed Molm14 cell lines*
Molm14 cells overexpressing WEE1 kinase domain-HiBiT were generated as previously reported (PMID: 37567174). Briefly, a codon-optimized gBlock (IDT) coding for amino acid residues 290-576 of WEE1 (Uniprot ID: P30291) with a C-terminal GSGS linker and HiBiT tag (VSGWRLFKKIS) was cloned by Gibson assembly for lentivirus-mediated expression under the control of the human EF1A promoter. To generate the WEE1 G322A mutant, QuickChange site-directed mutagenesis kit (Agilent; 200519) was used with the primer sequence listed below. Lentiviral transduction of Molm14 cells was carried out as previously reported (PMID: 37567174). Transduced Molm14 cells were grown under selection with 0.5 µg/mL of puromycin.

G322A_fwd: 5’-GTGTGAAGAGGCTGGATGCATGCATTTATGCCATTAAG-3’
G322A_rev: 5’-CTTAATGGCATAAATGCATGCATCCAGCCTCTTCACAC-3’

*HiBiT assays*

2.0 x $10^4$ HiBiT-tagged Jurkat cells were plated in 50 µL per well growth medium in opaque white 384-well plates (Corning). A D300 drug printer (Tecan) was used to dispense compounds from DMSO stock solutions at the indicated final concentrations. Cells were incubated at 37 ℃ for 5 hours. The Nano-Glo HiBiT Lytic Detection System (Promega) was used to measure endogenous indicated protein levels according to the manufacturer's instructions. All experiments (triplicates) were carried out at least twice on separate days. $DC_{50}$ values were determined using a non-linear regression curve fit in GraphPad PRISM 10.2.2 by considering each set of singlicate measurements in a given experiment as an independent replicate for curve fitting. The $DC_{50}$ is reported as the mean of the $DC_{50}$ values from the singlicate fits. The error was determined by calculating the standard error of the mean of the associated singlicate $DC_{50}$ values from the singlicate fits.

*Cell viability assays (CellTiter-Glo assay)*

2.0 x $10^3$ suspension cells were plated in 50 µL per well growth medium in opaque white 384-well plates (Corning) followed by drug treatment immediately at indicated concentrations. 1.0 x $10^3$ adherent cells were plated in 50 µL per well growth medium in opaque white 384-well plates (Corning) followed by drug treatment the next day at indicated concentrations. After 72 hours incubation, cellular ATP content was measured using CellTiter-Glo reagent (Promega). Cell viability and $IC_{50}$ values were determined using a non-linear regression curve fit in GraphPad PRISM 10.2.2 by considering each set of singlicate measurements in a given experiment as an independent replicate for curve fitting. The $IC_{50}$ is reported as the mean of the $DC_{50}$ values from the singlicate fits. The error was determined by calculating the standard error of the mean of the associated singlicate $DC_{50}$ values from the singlicate fits.

*Construct and protein purification*

Codon-optimized WEE1 kinase domain (residues 291-575; Uniprot ID 30291-1) was cloned by ligation-independent cloning into a vector coding for an N-terminal His6-GST-TEVsite tag.

The resulting plasmid was grown in Rosetta2(DE3)pLysS chemically competent *E. coli* cells (Novagene) and grown in Luria Broth supplemented with chloramphenicol and carbenicillin overnight. Saturated overnight cultures were distributed to 1 L flasks containing 2XYT medium supplemented with antibiotics. Protein expression was induced by addition of 400 µM IPTG (final concentration). The temperature was adjusted from 37°C to 18°C. After incubation overnight, cells were harvested by centrifugation. Cell pellets were resuspended in ~4 ml/L D800 buffer (20 mM HEPES, pH 7.5; 800 mM NaCl; 10 mM imidazole, pH 8.0; 10 % glycerol, 2 mM beta mercaptoethanol) supplemented with protease inhibitors (1 mM PMSF, 1 mM benzamidine, ~20 ug/ml pepstatin, aprotinin, and leupeptin) and frozen at -80°C.

Protein purification was carried out essentially as described previously.[1] Frozen *E. coli* cell pellets were thawed, sonicated, and centrifuged for one hour at 12 degrees Celsius. The resulting soluble supernatant was incubated with cobalt agarose beads for one hour at 4 degrees Celsius, after which the beads were washed with ~50 column volumes D800 followed by a single wash in B50 buffer (D800 with 50 mM NaCl). Recombinant kinase was eluted from the beads by several washes with buffer C50 (B50 with 400 mM imidazole, pH 8.0). The eluate was applied to an anion exchange column (Q HP; Cytiva) equilibrated in B50 and eluted by a linear gradient to D800. Peak fractions were pooled, concentrated by ultrafiltration, and applied to a 24 ml gel filtration column (S200 increase, Cytiva) primed with GF150 buffer (20 mM Tris-HCl, pH 8.5, 150 mM NaCl, 1 mM TCEP). Peak fractions were again concentrated by ultrafiltration, supplemented with 5% glycerol (v:v, final), and aliquoted and frozen at -80°C.

GST-TEV-CRBN and DDB1 were cloned into pLIB vectors. Constructs were transformed into DH10EmBacY cells, and the derived bacmids were transfected with FuGENE HD into *Trichoplusia Ni* Sf9 insect cells for baculovirus generation. Hi5 insect cells were coinfected with GST-TEV-CRBN and DDB1 viruses for expression of the CRBN-DDB1 complex. Proteins were purified through GST affinity purification, followed by an overnight TEV cleavage, IEX by PorosHQ, and size exclusion chromatography into a 25mM HEPES pH 7.5, 150mM NaCl, 1mM TCEP. WEE1 protein used for cryo-EM was prepared essentially as described above, except N-terminal affinity tags were removed after the ion exchange step by incubation with TEV protease for three hours before removal of the protease, cleaved tags, and uncleaved WEE1 by nickle affinity chromatography. Untagged WEE1 was concentrated and subjected to size exclusion chromatography before further concentration and storage as described above. CRBN-DDB1 was purified with the Flag tag, Spycatcher was conjugated with BODIPY FL, and the two were combined to make the BODIPY-CRBN-DDB1 following a previously reported procedure.[2]

*Cryo-EM sample preparation and data collection*

Complex was formed at 10µM CRBN-DDB1, 15µM WEE1, and 100µM compound 10 for 30 minutes on ice in 25mM HEPES pH 7.5, 150mM NaCl, 1mM TCEP. The complex was diluted immediately prior to vitrification at various concentrations between 1-2.5µM suitable for optimal particle distribution. Quantifoil UltrAuFoil R1.2/1.3 300 mesh grids were glow discharged in PELCO easiglow at 20mA for 2min. Glow-charged grids were preincubated with 4µL of 10µM CRBN agnostic IKZF1 ZF2 Q146A G151N as described[3] for 1min. After incubation, grid was blotted from the back, followed by 4µL CRBN complex sample application and vitrification into liquid ethane by a Leica EM-GP.

Grids were screened and collected on a Talos Arctica with a Gatan K3 direct electron detector. Data was collected at 200kV, 1.1 Å/pix with a nominal magnification at 36,000X, a total electron dose of 53e-/Å$^2$ in 50 frames, defocus range of -1.0 ~ -2.2µm. 3229 final accepted movies after filtering through micrograph quality were collected and processed using cryoSPARC live,[4]

2,766,285 particles were picked with a 2D template. After iterative 2D classifications, 3D classifications, and various 3D refinements, 101,857 particles were used for the final refinement. DeepEMhancer[5] was used for the map post-processing and map resolutions were estimated based on a FSC threshold at 0.143. Structural analyses were done by docking published structures of CRBN-DDB1 (PDB ID: 5FQD),[6] and WEE1 (PDB ID: 8BJU)[7] using softwares including Chimera,[8] ChimeraX,[9] Pymol, and Coot.[10]

| Data collection and processing | WEE1-compound 10-CRBN-DDB1 CRBN closed conformation EMD-XXXXX | WEE1-compound 10-CRBN-DDB1 CRBN open conformation EMD-XXXXX |
|---|---|---|
| Microscope | Talos Arctica | |
| Magnification | 36000 | |
| Voltage (kV) | 200 | |
| Electron exposure (e–/Å$^2$) | 53 | |
| Defocus range (µm) | -1.0 ~ -2.2 | |
| Automation software | SerialEM | |
| Pixel size (Å) | 1.1 | |
| Symmetry imposed | C1 | |
| Movies collected | 3229 | |
| Initial particle (no.) | 2,766,285 | |
| Final particle (no.) | 101,857 | 87,492 |
| Map resolution(Å)<br>FSC threshold | 3.73<br>(0.143) | 4.13<br>(0.143) |

*Time-Resolved Fluorescence Resonance Energy Transfer Assay*

Compounds in the binding assay were dispensed into a 384-well plate (Corning, cat#4514) containing the assay mix (10 uL/well) using the D300 drug printer (Tecan) normalized to 1% DMSO. Assay mix composed of 100 nM WEE1 (His-GST), 10 nM Flag-Spy(BODIPY FL labeled)-CRBN + His-DDB1ΔBPB and diluted (1:400) Europium-labeled anti-6X His antibody (PerkinElmer, cat#AD0402) prepared in assay buffer with 20 mM HEPES, 150 mM NaCl, 1mM fresh TCEP, 0.1% BSA, and 0.1% NP-40 as described previously.[2] The reactions were incubated for 2 hours at room temperature in dark before TR-FRET measurements were conducted. After excitation of europium fluorescence at 337 nm, emission at 490 nm (europium) and 520 nm (Alexa Fluor 647) were recorded, with a 70 µs delay over 600 µs to reduce background fluorescence, using a PHERAstar FSX plate reader (BMG Labtech) as described previously.[11] The TR-FRET signal was calculated as the 490/520 nm ratio for each time point and plotted by GraphPad Prism 7.

*Global quantitative proteomics sample preparation*

Cells were lysed by addition of lysis buffer (8 M Urea, 50 mM NaCl, 50 mM 4-(2-hydroxyethyl)-1-piperazineethanesulfonic acid (EPPS) pH 8.5, Protease and Phosphatase inhibitors) and homogenization by bead beating (BioSpec) for three repeats of 30 seconds at 2400 strokes/min. Bradford assay was used to determine the final protein concentration in the clarified cell lysate. Fifty micrograms of protein for each sample was reduced, alkylated and precipitated using methanol/chloroform as previously described[12] and the resulting washed precipitated protein was allowed to air dry. Precipitated protein was resuspended in 4 M urea, 50 mM HEPES pH 7.4, followed by dilution to 1 M urea with the addition of 200 mM EPPS, pH 8. Proteins were digested with the addition of LysC (1:50; enzyme:protein) and trypsin (1:50; enzyme:protein) for 12 h at 37 °C. Sample digests were acidified with formic acid to a pH of 2-3 before desalting using C18 solid phase extraction plates (SOLA, Thermo Fisher Scientific). Desalted peptides were dried in a vacuum-centrifuged and reconstituted in 0.1% formic acid for liquid chromatography-mass spectrometry analysis.

Data were collected using a TimsTOF Pro2 (Bruker Daltonics, Bremen, Germany) coupled to a nanoElute LC pump (Bruker Daltonics, Bremen, Germany) via a CaptiveSpray nano-electrospray source. Peptides were separated on a reversed-phase $C_{18}$ column (25 cm x 75 µm ID, 1.6 µM, IonOpticks, Australia) containing an integrated captive spray emitter. Peptides were separated using a 50 min gradient of 2 - 30% buffer B (acetonitrile in 0.1% formic acid) with a flow rate of 250 nL/min and column temperature maintained at 50 ºC.

Data-dependent acquisition (DDA) was performed in parallel accumulation-serial fragmentation (PASEF) mode to determine effective ion mobility windows for downstream diaPASEF data collection.[13] The diaPASEF parameters included: 100% duty cycle using accumulation and ramp times of 50 ms each, 1 TIMS-MS scan and 10 PASEF ramps per acquisition cycle. The TIMS-MS survey scan was acquired between 100 – 1700 *m/z* and 1/k0 of 0.7 - 1.3 V.s/cm$^2$. Precursors with 1 – 5 charges were selected and those that reached an intensity threshold of 20,000 arbitrary units were actively excluded for 0.4 min. The quadrupole isolation width was set to 2 *m/z* for *m/z* <700 and 3 *m/z* for *m/z* >800, with the *m/z* between 700-800 *m/z* being interpolated linearly. The TIMS elution voltages were calibrated linearly with three points (Agilent ESI-L Tuning Mix Ions; 622, 922, 1,222 *m/z*) to determine the reduced ion mobility coefficients ($1/K_0$). To perform diaPASEF, the precursor distribution in the DDA *m/z*-ion mobility plane was used to design an acquisition scheme for Data-independent acquisition (DIA) data collection which included two windows in each 50 ms diaPASEF scan. Data was acquired using sixteen of these 25 Da precursor double window scans (creating 32 windows) which covered the diagonal scan line for doubly and triply charged precursors, with singly charged precursors able to be excluded by their position in the m/z-ion mobility plane. These precursor isolation windows were defined between 400 - 1200 *m/z* and 1/k0 of 0.7 - 1.3 V.s/cm$^2$.
