## Supplementary material for "Discovery of CRBN-dependent WEE1 Molecular Glue Degraders from a Multicomponent Combinatorial Library": Synthetic Methods

**Synthetic Methods for:**

**General synthetic methods**

Unless otherwise noted, all reagents were purchased from commercial suppliers and used without further purification. Reactions were monitored using a Waters Acquity UPLC/MS system (Waters PDA eλ Detector, QDa Detector, Sample manager - FL, Binary Solvent Manager) using Acquity UPLC® BEH C18 column (2.1 x 50 mm, 1.7 μm particle size): solvent gradient = 85% A at 0 min, 1% A at 1.7 min; solvent A = 0.1% formic acid in water; solvent B = 0.1% formic acid in Acetonitrile; flow rate: 0.6 mL/min. Analytical thin layer chromatography (TLC) was performed on Merck silica gel 60 F254 TLC glass plates and analytes were visualized by fluorescence quenching (using 254 nm light). Purification of reaction products was carried out by flash column chromatography using CombiFlash®Rf with Teledyne Isco RediSep® normal-phase silica flash columns (4 g, 12 g, 24 g, 40 g or 80 g) or preparative RP-HPLC using Waters SunFireTM Prep C18 column (19 x 100 mm, 5 μm particle size) using a gradient of 5-95% methanol in water containing 0.05% trifluoroacetic acid (TFA) over 40 min (45 min run time) at a flow of 40 mL/min. Assayed compounds were isolated and tested as TFA salts and purities of assayed compounds were in all cases greater than 95%, as determined by reverse-phase UPLC analysis. NMR spectra were acquired on a 500 MHz Bruker Avance III spectrometer, operating at the denoted spectrometer frequency given in MHz for the specified nucleus. All experiments were acquired at 298.0 K with a calibrated Bruker Variable Temperature Controller unless otherwise noted. The chemical shifts are reported in parts per million (ppm) and coupling constants (J) are given in Hertz (Hz). 1H NMR spectra are reported with the solvent resonance as the reference unless noted otherwise (DMSO-d6 at 2.50 ppm). Peaks are reported as (s = singlet, d = doublet, t = triplet, q = quartet, m = multiplet or unresolved, br = broad signal, coupling constant(s) in Hz, integration).

### Compound Synthesis

**Scheme 1.** Synthesis of the HRZ-1 scaffold aldehyde precursor (compound D)

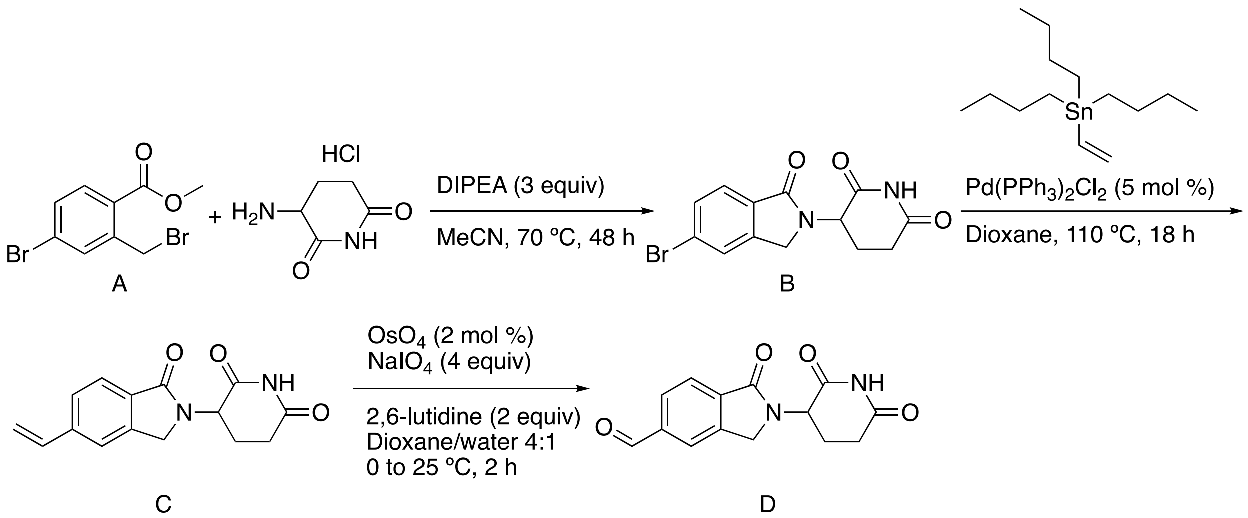


**Compound B**. Synthesized following a reference procedure.[1] To the solution of A (1.0 g, 3.2 mmol, 1.0 equiv) and 3-aminopiperidine-2,6-dione hydrochloride (0.80 g, 4.9 mmol, 1.5 equiv) in acetonitrile (12 mL) was added DIPEA (1.7 mL, 9.7 mmol, 3.0 equiv) under nitrogen flow. The resulting suspension was stirred at 80 °C for 48 hours. The reaction mixture was cooled to 4 ºC, upon which a precipitate formed. The solid was filtered off, washed with cold MeCN (30 mL), MeCN:Et2O (25 mL [2:3]), and $Et_2O$ (2 x 25 mL) to afford compound B as a dark blue solid that was used directly in the next step (0.79 g, 76%). **LC-MS**: calculated exact mass = 322.00; found $[M+H]^+$ = 323.06.

**Compound C**. Synthesized following a reference procedure.[2] To a solution of compound B ( 0.79 g, 2.5 mmol, 1.0 equiv) in 1,4-Dioxane (30 mL) was added tributyl(vinyl)stannane (1.1 mL, 3.7 mmol, 1.5 equiv) and Bis-(triphenylphosphino)-palladous chloride (86 mg, 0.12 mmol, 0.05 equiv) and the mixture was stirred at 110 °C for 18 hour. Confirmed product formation by LC-MS and diluted with water and extracted with EtOAc 3 times. The combined organic layer was washed with brine, dried over sodium sulfate, and concentrated under reduced pressure onto a silica pad. The compound C was isolated via silica gel chromatography in hexane:EtOAc gradient elution and used directly in the next step (530 mg, 80%). **LC-MS**: calculated exact mass = 270.10; found $[M+H]^+$ = 271.24.

**Compound D**. Synthesized following a reference procedure.[2] To a solution of compound C (0.53 g, 2.0 mmol, 1.0 equiv) in Water (5.0 mL) and 1,4-dioxane (20 mL) was added sodium periodate (1.7 g, 7.8 mmol, 4.0 equiv), 2,6-dimethylpyridine (0.45 mL, 3.9 mmol, 2.0 equiv), and the mixture was cooled to 0 ºC. Osmium(VIII) oxide (0.24 mL, 4% w/v, 0.039 mmol, 0.02 equiv) was added dropwise and the mixture was allowed to warm up to 25 °C after 10 min. The color changed from deep blue to colorless and a white precipitate formation was observed. The mixture was stirred for 2 hour and then diluted with 20 mL of water, extracted 3 x 15 mL of EtOAc. Organic layer was washed with brine, dried, and evaporated. The compound D was isolated as a white solid via silica gel chromatography using hexane:EtOAc gradient elution (790 mg, 76%). **LC-MS**: calculated exact mass = 272.08; found $[M+H]^+$ = 273.14. **$^1$H NMR** (500 MHz, DMSO-$d_6$) δ = 11.02 (s, 1H), 10.15 (s, 1H), 8.15 (s, 1H), 8.06 (dd, *J* = 7.8, 1.4 Hz, 1H), 7.94 (d, *J* = 7.8 Hz, 1H), 5.16 (dd, *J* = 13.3, 5.1 Hz, 1H), 4.58 (d, *J* = 17.7 Hz, 1H), 4.45 (d, *J* = 17.7 Hz, 1H), 2.92 (m, 1H), 2.68 – 2.56 (m, 1H), 2.42 (m, 1H), 2.13 – 1.99 (m, 1H).

**Compound E**

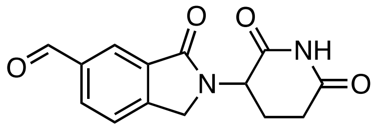


Compound E was synthesized following the synthetic route in Scheme 1 starting from 3-(6-bromo-1-oxoisoindolin-2-yl)piperidine-2,6-dione in place of compound B. Yield: 240 mg, 31% over two steps. **LC-MS**: calculated exact mass = 272.08; found $[M+H]^+$ = 273.24.

**Compound F**

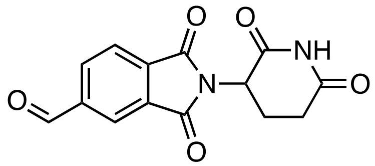


Compound F was synthesized following the synthetic route in Scheme 1 starting from 3-(6-bromo-1-oxoisoindolin-2-yl)piperidine-2,6-dione in place of compound B. Yield: 850 mg, 50% over two steps. **LC-MS**: calculated exact mass = 286.06; found $[M-H]^-$ = 285.09.

**General procedures for Groebke-Blackburn-Bienaymé (GBB) cyclization**

**Procedure A**. Aldehyde (1.1 equiv), amidine (1.0 equiv), and acid catalyst were dissolved in anhydrous MeOH (200-300 mM final aldehyde molarity) and stirred for 30-60 minutes. Isocyanide (1.1 equiv) was then added, and the reaction was stirred for 24-72 h. The product formation was detected by LC-MS and/or TLC. The mixture was then diluted in DMSO and purified via preparative HPLC using MeOH:$H_2O$ (0.035% formic acid additive) elution gradient.

**Procedure B.** Stock solutions of aldehyde (1.0 equiv per reaction) in NMP (solution A) and amidine (1.1 equiv per reaction) with acid catalyst in MeOH (solution B) were prepared. Mixed specified volumes of A and B to reach 150-300 mM final amidine concentration and desired aldehyde:amidine stoichiometry. After stirring for 30-60 minutes, isocyanide (1.2 equiv) was added, and the reaction was stirred for 24-72 h. The product formation was monitored by LC-MS and/or TLC. The mixture was then diluted in DMSO and purified via preparative HPLC using MeOH:$H_2O$ (0.035% formic acid additive) elution gradient.

**Compound 1**

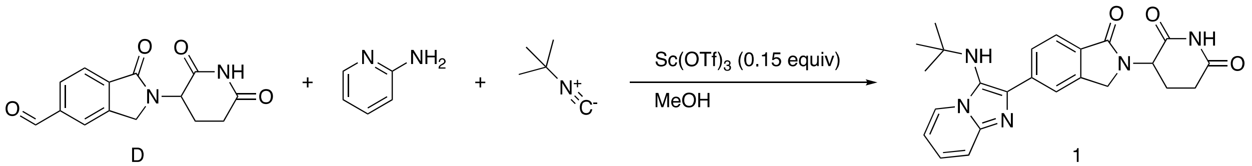


D 1

Compound 1 was synthesized following procedure A. Compound D was used as the aldehyde precursor along with 0.15 equiv of $Sc(OTf)_3$ as the acid catalyst. Yield: 14 mg, 43%. **LC-MS**: calculated exact mass = 431.20; found $[M+H]^+$ = 432.35. **$^1$H NMR** (500 MHz, DMSO-$d_6$) δ = 11.03 (s, 1H), 8.86 (d, *J* = 6.8 Hz, 1H), 8.28 (s, 1H), 8.22 (d, *J* = 7.9, 1H), 7.93 (d, *J* = 7.9 Hz, 1H), 7.87 (s, 2H), 7.46 (s, 1H), 5.24 (s, 1H), 5.16 (dd, *J* = 13.3, 5.1 Hz, 1H), 4.59 (d, *J* = 17.4 Hz, 1H), 4.46 (d, *J* = 17.4 Hz, 1H), 2.94 (ddd, *J* = 17.3, 13.6, 5.4 Hz, 1H), 2.67 – 2.58 (m, 1H), 2.44 (m, 1H), 2.07 (m, 1H), 1.01 (s, 9H).

**Compound 2**

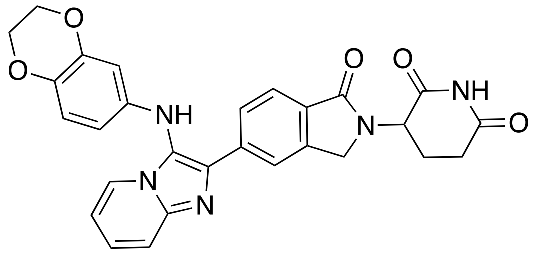

Compound 2 was synthesized following procedure A. Compound D was used as the aldehyde precursor along with 0.15 equiv of $Sc(OTf)_3$ as the acid catalyst. Yield: 16 mg, 42%. **LC-MS**: calculated exact mass = 509.17; found $[M+H]^+$ = 510.22. **$^1$H NMR** (500 MHz, DMSO-$d_6$) δ = 11.00 (s, 1H), 8.27 (s, 1H), 8.23 – 8.17 (m, 2H), 8.12 (d, *J* = 8.0 Hz, 1H), 7.86 (d, *J* = 8.1 Hz, 2H), 7.72 (s, 1H), 7.27 (s, 1H), 6.71 – 6.64 (m, 1H), 6.19 – 6.11 (m, 2H), 5.14 (dd, *J* = 13.3, 5.1 Hz, 1H), 4.53 (d, *J* = 17.4 Hz, 1H), 4.39 (d, *J* = 17.4 Hz, 1H), 4.16 – 4.13 (m, 2H), 4.13 – 4.11 (m, 2H), 2.92 (ddd, *J* = 16.9, 13.4, 5.2 Hz, 1H), 2.65 – 2.56 (m, 1H), 2.48 – 2.36 (m, 1H), 2.05 – 1.98 (m, 1H).

**Compound 3**

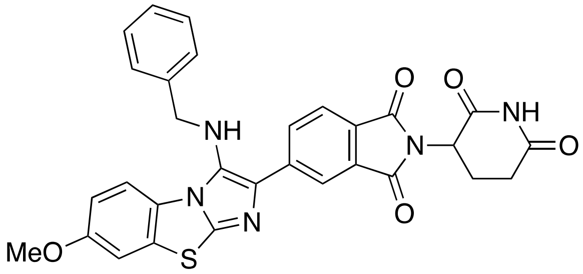

Compound 3 was synthesized following procedure B. Compound D was used as the aldehyde precursor (1:1.1 ratio to amidine) along with 0.15 equiv of $Sc(OTf)_3$ as the acid catalyst. 200 µL of solution A and 50 µL of solution B were used. Yield: 2.0 mg, 7% **LC-MS**: calculated exact mass = 551.16 found $[M+H]^+$ = 552.39. **$^1$H NMR** (500 MHz, DMSO-$d_6$) δ = 11.00 (s, 1H), 8.18 – 8.07 (m, 2H), 8.00 (d, *J* = 8.9 Hz, 1H), 7.69 (d, *J* = 7.9 Hz, 1H), 7.64 (d, *J* = 2.6 Hz, 1H), 7.32 (d, *J* = 6.9 Hz, 2H), 7.27 (t, *J* = 7.6 Hz, 2H), 7.23 – 7.18 (m, 1H), 7.09 (dd, *J* = 9.0, 2.6 Hz, 1H), 5.49 (s, 1H), 5.13 (dd, *J* = 13.3, 5.1 Hz, 1H), 4.45 (d, *J* = 17.0 Hz, 1H), 4.33 (d, *J* = 17.0 Hz, 1H), 4.18 (s, 2H), 3.83 (s, 3H), 2.93 (ddd, *J* = 17.2, 13.6, 5.4 Hz, 1H), 2.65 – 2.58 (m, 1H), 2.48 – 2.37 (m, 1H), 2.07 – 1.99 (m, 1H).

**Compound 4**

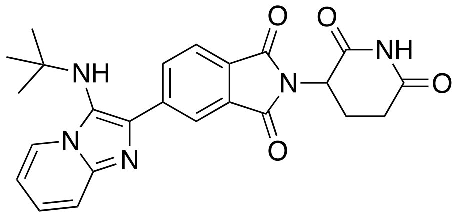

Compound 4 was synthesized following procedure B. Compound F was used as the aldehyde precursor along with 0.15 equiv of $Sc(OTf)_3$ as the acid catalyst. 200 µL of solution A and 150 µL of solution B were used. Yield: 12 mg, 39%. **LC-MS**: calculated exact mass = 445.18; found $[M+H]^+$ = 446.35. $^1$**H NMR** (500 MHz, DMSO-$d_6$) δ = 11.16 (s, 1H), 8.77 (d, *J* = 6.9 Hz, 1H), 8.65 (s, 1H), 8.62 (dd, *J* = 7.8, 1.5 Hz, 1H), 8.10 (d, *J* = 7.8 Hz, 1H), 7.82 (d, *J* = 8.9 Hz, 1H), 7.79 – 7.73 (m, 1H), 7.36 (t, *J* = 6.8 Hz, 1H), 5.28 (s, 1H), 5.20 (dd, *J* = 12.8, 5.5 Hz, 1H), 2.91 (ddd, *J* = 16.8, 13.7, 5.4 Hz, 1H), 2.70 – 2.53 (m, 2H), 2.18 – 2.07 (m, 1H), 1.04 (s, 9H).

**Compound 5**

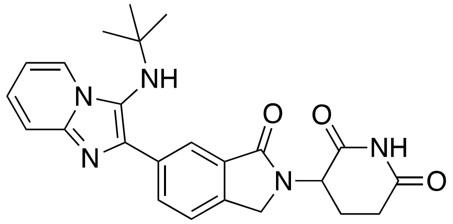

Compound 5 was synthesized following procedure B. Compound E was used as the aldehyde precursor along with 0.15 equiv of $Sc(OTf)_3$ as the acid catalyst. 200 µL of solution A and 150 µL of solution B were used. Yield: 6.2 mg, 21%. **LC-MS**: calculated exact mass = 431.20; found $[M+H]^+$ = 432.40. $^1$**H NMR** (500 MHz, DMSO-$d_6$) δ = 11.03 (s, 1H), 8.84 (d, *J* = 6.9 Hz, 1H), 8.42 (s, 1H), 8.37 (dd, *J* = 8.0, 1.7 Hz, 1H), 7.86 (s, 2H), 7.82 (d, *J* = 8.0 Hz, 1H), 7.50 – 7.41 (m, 1H), 5.26 (s, 1H), 5.16 (dd, *J* = 13.3, 5.1 Hz, 1H), 4.58 (d, *J* = 17.8 Hz, 1H), 4.44 (d, *J* = 17.7 Hz, 1H), 2.93 (ddd, *J* = 17.3, 13.6, 5.4 Hz, 1H), 2.67 – 2.58 (m, 1H), 2.48 – 2.39 (m, 1H), 2.10 – 2.01 (m, 1H), 1.02 (s, 9H).

**Compound 10**

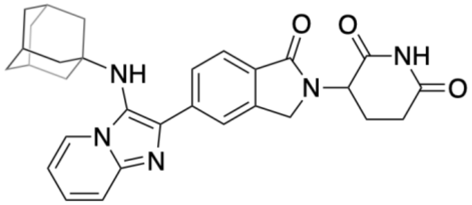

Compound 10 was synthesized following procedure B. Compound D was used as the aldehyde precursor along with 0.15 equiv of $Sc(OTf)_3$ as the acid catalyst. 200 µL of solution A and 150 µL of solution B were used. Yield: 27.3 mg, 72%. **LC-MS**: calculated exact mass = 509.24; found $[M+H]^+$ = 510.42. **$^1$H NMR** (500 MHz, DMSO-$d_6$) δ = 11.03 (s, 1H), 8.87 (d, *J* = 6.8 Hz, 1H), 8.34 – 8.21 (m, 2H), 7.92 (d, *J* = 7.9 Hz, 1H), 7.86 (d, *J* = 2.2 Hz, 2H), 7.50 – 7.36 (m, 1H), 5.25 (s, 1H), 5.16 (dd, *J* = 13.3, 5.2 Hz, 1H), 4.60 (d, *J* = 17.5 Hz, 1H), 4.46 (d, *J* = 17.5 Hz, 1H), 2.94 (ddd, *J* = 17.3, 13.7, 5.4 Hz, 1H), 2.66 – 2.59 (m, 1H), 2.44 (td, *J* = 13.2, 4.5 Hz, 1H), 2.11 – 2.02 (m, 1H), 1.91 – 1.86 (m, 3H), 1.58 (d, *J* = 2.9 Hz, 6H), 1.52 – 1.39 (m, 6H).

**Compound 11**

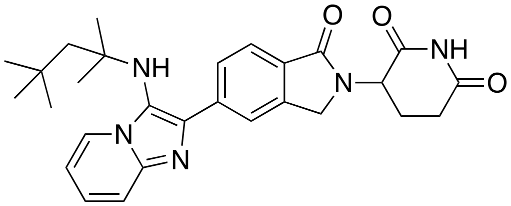

Compound 11 was synthesized following procedure B. Compound D was used as the aldehyde precursor along with 0.15 equiv of $Sc(OTf)_3$ as the acid catalyst. 250 µL of solution A and B were used. Yield: 27 mg, 81%. **LC-MS**: calculated exact mass = 487.26; found $[M+H]^+$ = 488.42. **$^1$H NMR** (500 MHz, DMSO-$d_6$) δ = 11.02 (s, 1H), 8.87 (d, *J* = 6.8 Hz, 1H), 8.23 (s, 1H), 8.15 (dd, *J* = 8.0, 1.4 Hz, 1H), 7.96 – 7.86 (m, 3H), 7.50 (t, *J* = 6.7 Hz, 1H), 5.17 (dd, *J* = 13.3, 5.2 Hz, 1H), 5.05 (s, 1H), 4.58 (d, *J* = 17.5 Hz, 1H), 4.46 (d, *J* = 17.5 Hz, 1H), 2.94 (ddd, *J* = 17.3, 13.6, 5.4 Hz, 1H), 2.66 – 2.59 (m, 1H), 2.48 – 2.41 (m, 1H), 2.10 – 2.02 (m, 1H), 1.58 (s, 2H), 0.98 – 0.91 (m, 15H).

**Compound 12**

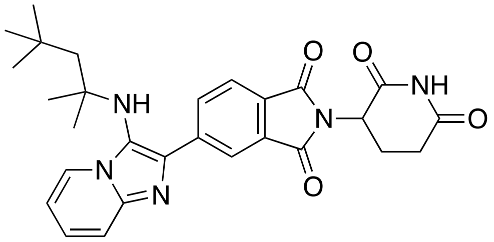

Compound 12 was synthesized following procedure B. Compound F was used as the aldehyde precursor (1:1.2 ratio to amidine) along with 0.15 equiv of $Sc(OTf)_3$ as the acid catalyst. 150 µL of solution A and 50 µL of solution B were used. Yield: 18 mg, 59% **LC-MS**: calculated exact mass = 501.24 found $[M+H]^+$ = 502.32. **$^1$H NMR** (500 MHz, DMSO-$d_6$) δ = 11.16 (s, 1H), 8.77 (d, *J* = 6.9 Hz, 1H), 8.64 (s, 1H), 8.55 (dd, *J* = 7.8, 1.5 Hz, 1H), 8.11 (d, *J* = 7.9 Hz, 1H), 7.82 (d, *J* = 8.8 Hz, 1H), 7.79 – 7.73 (m, 1H), 7.40 – 7.33 (m, 1H), 5.20 (dd, *J* = 12.8, 5.4 Hz, 1H), 5.07 (s, 1H), 2.91 (ddd, *J* = 16.6, 13.7, 5.4 Hz, 1H), 2.66 – 2.60 (m, 1H), 2.59 – 2.53 (m, 1H), 2.14 – 2.07 (m, 1H), 1.63 (s, 2H), 0.99 (s, 9H), 0.96 (s, 6H).

**Compound 13**

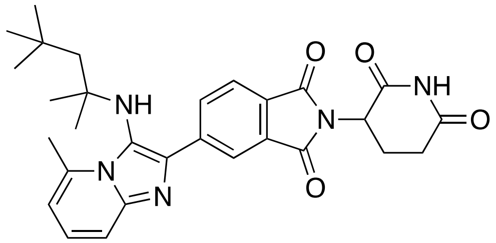

Compound 13 was synthesized following procedure B. Compound F was used as the aldehyde precursor (1:1.2 ratio to amidine) along with 0.15 equiv of $Sc(OTf)_3$ as the acid catalyst. 150 µL of solution A and 50 µL of solution B were used. Yield: 8.0 mg, 25% **LC-MS**: calculated exact mass = 515.25 found $[M+H]^+$ = 516.33. **$^1$H NMR** (500 MHz, DMSO-$d_6$) δ = 11.16 (s, 1H), 8.54 (s, 1H), 8.46 (dd, *J* = 7.8, 1.5 Hz, 1H), 8.12 (d, *J* = 7.8 Hz, 1H), 7.70 – 7.65 (m, 2H), 7.10 (s, 1H), 5.21 (dd, *J* = 12.9, 5.4 Hz, 1H), 4.87 (s, 1H), 3.08 (s, 3H), 2.91 (ddd, *J* = 16.6, 13.6, 5.3 Hz, 1H), 2.66 – 2.61 (m, 1H), 2.60 – 2.53 (m, 1H), 2.16 – 2.07 (m, 1H), 1.40 (s, 2H), 0.86 (s, 9H), 0.83 (s, 6H).

**Compound 1-Me**

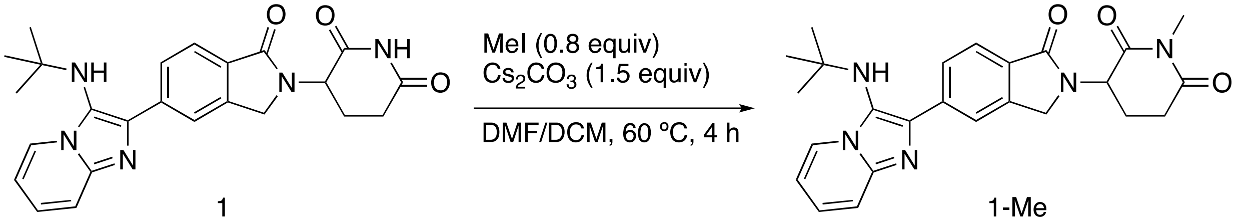


To the solution of compound 1 (7.0 mg, 16 µmol, 1.0 equiv) in 200 µL DMF was added $Cs_2CO_3$ (7.9 mg, 24 µmol, 1.5 equiv). Then MeI (13 µL, 1 M in DCM, 13 µmol, 0.8 equiv) were added and the mixture was stirred at 60 ºC for 4 hours. LCMS analysis confirmed single methylation product and the mixture was diluted with DMSO, filtered, and purified via preparative HPLC using MeOH:$H_2O$ gradient. Yield: 3.7 mg, 64%. **LC-MS**: calculated exact mass = 445.21 found $[M+H]^+$ = 446.30. **$^1$H NMR** (500 MHz, DMSO-$d_6$) δ = 8.85 (d, *J* = 6.9 Hz, 1H), 8.33 (s, 1H), 8.27 (d, *J* = 1.4 Hz, 1H), 7.94 – 7.86 (m, 3H), 7.44 (s, 1H), 5.28 – 5.19 (m, 2H), 4.58 (d, *J* = 17.4 Hz, 1H), 4.44 (d, *J* = 17.4 Hz, 1H), 3.07 – 2.96 (m, 4H), 2.83 – 2.74 (m, 1H), 2.49 – 2.39 (m, 1H), 2.12 – 2.03 (m, 1H), 1.02 (s, 9H).

**Compound 6**

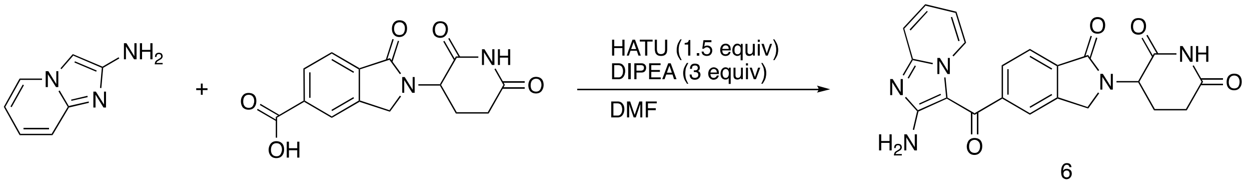


To a solution of 2-(2,6-dioxopiperidin-3-yl)-1-oxoisoindoline-5-carboxylic acid (20 mg, 0.070 mmol) and DIPEA (36 µL, 0.21 mmol, 3.0 equiv) in DMF (0.3 mL) was added HATU (40 mg, 0.10 mmol, 1.5 equiv) and imidazo[1,2-a]pyridin-2-amine (11mg, 0.080 mmol, 1.1 equiv) at room temperature. The mixture was stirred at room temperature for 1 h, and then quenched by sodium bicarbonate sat. solution, extracted by Chloroform/isopropanol (4/1), concentrated *in vacuo*. The residue was dissolved in DMSO and purified via preparative HPLC using MeOH:$H_2O$ gradient. Yield: 5.7 mg, 21%. **LC-MS**: calculated exact mass = 403.13 found $[M+H]^+$ = 404.23. **$^1$H NMR** (500 MHz, DMSO-$d_6$) δ = 11.02 (s, 1H), 9.18 (s, 1H), 7.88 (d, *J* = 7.7 Hz, 1H), 7.83 (s, 1H), 7.71 (dd, *J* = 7.8, 1.4 Hz, 1H), 7.59 (ddd, *J* = 8.6, 7.0, 1.3 Hz, 1H), 7.43 (d, *J* = 8.5 Hz, 1H), 7.05 (td, *J* = 6.9, 1.3 Hz, 1H), 5.96 (s, 2H), 5.17 (dd, *J* = 13.3, 5.1 Hz, 1H), 4.54 (d, *J* = 17.6 Hz,

1H), 4.43 (d, $J$ = 17.5 Hz, 1H), 2.93 (ddd, $J$ = 17.3, 13.6, 5.4 Hz, 1H), 2.65 – 2.57 (m, 1H), 2.49 – 2.36 (m, 1H), 2.10 – 1.99 (m, 1H).

**Compound 7**

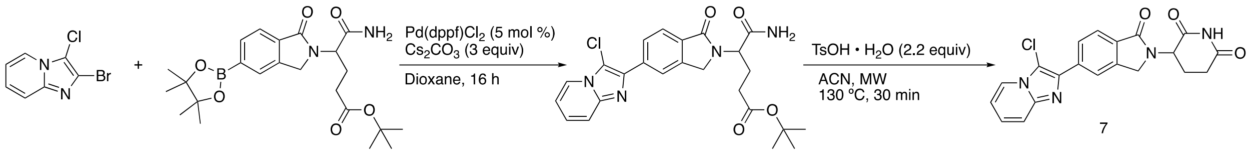


5:3 dioxane:water solvent mixture was degassed under continuous nitrogen flow for at least 10 minutes. A vial was charged with 2-bromo-3-chloroimidazo[1,2-a]pyridine (14 mg, 62 µmol, 1.1 equiv), tert-butyl 5-amino-5-oxo-4-(1-oxo-5-(4,4,5,5-tetramethyl-1,3,2-dioxaborolan-2-yl)isoindolin-2-yl)pentanoate (25 mg, 56 µmol, 1.0 equiv), Pd(dppf)$Cl_2$ • DCM (2.3 mg, 2.8 µmol, 0.05 equiv) and $K_3PO_4$ (60 mg, 0.28 mmol, 5.0 equiv). After three vacuum/nitrogen cycles, degassed solvent mixture was added and heated overnight at 70 °C. LCMS showed the coupling product formation with partial tert-butyl ester hydrolysis. The mixture was diluted with EtOAc, passed through wet celite pad, and dried under vacuum. The solid residue was dissolved in ACN (500 µL) and TsOH•$H_2O$ (24 mg, 120 µmol, 2.2 equiv) were added. The mixture was diluted with DMSO and purified via preparative HPLC using MeOH:$H_2O$ gradient. Yield (over two steps): 4.6 mg, 21%. **LC-MS**: calculated exact mass = 394.08 found $[M+H]^+$ = 395.13. $^1H$ NMR (500 MHz, DMSO) δ 11.01 (s, 1H), 8.45 (d, $J$ = 6.9 Hz, 1H), 8.33 (s, 1H), 8.26 (dd, $J$ = 7.9, 1.4 Hz, 1H), 7.88 (d, $J$ = 8.0 Hz, 1H), 7.72 (d, $J$ = 9.2 Hz, 1H), 7.45 (ddd, $J$ = 9.1, 6.8, 1.3 Hz, 1H), 7.17 (td, $J$ = 6.8, 1.1 Hz, 1H), 5.15 (dd, $J$ = 13.3, 5.2 Hz, 1H), 4.57 (d, $J$ = 17.3 Hz, 1H), 4.44 (d, $J$ = 17.3 Hz, 1H), 2.93 (ddd, $J$ = 17.4, 13.6, 5.4 Hz, 1H), 2.66 – 2.59 (m, 1H), 2.49 – 2.37 (m, 1H), 2.09 – 2.00 (m, 1H).

**Compound 8**

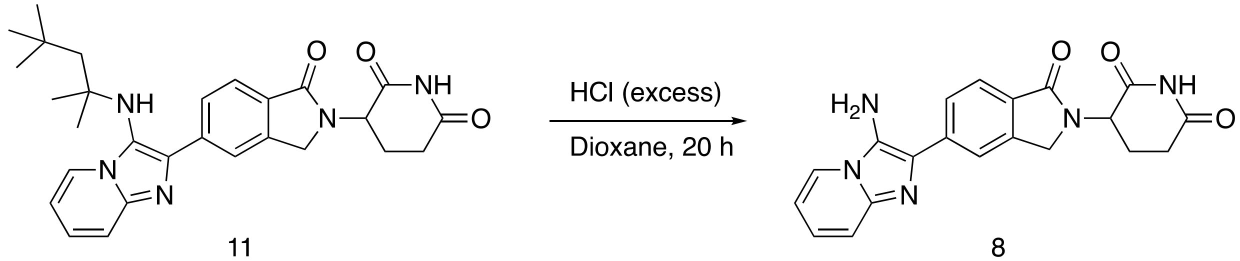


Compound 11 (0.49 g, 1.0 mmol, 1.0 equiv) was dissolved in dioxane (5 mL). 4 M HCl in dioxane (5 mL) was added and the mixture was stirred for 20 hours, until complete conversion was detected by LC-MS. The mixture was then diluted in DMSO and purified via preparative HPLC in portions using MeOH:$H_2O$ (0.035% formic acid additive) elution gradient. Yield: 280 mg, 75%. **LC-MS**: calculated exact mass = 375.13 found $[M+H]^+$ = 376.33. **$^1$H NMR** (500 MHz, DMSO-$d_6$) δ = 11.02 (s, 1H), 8.67 (d, *J* = 6.9 Hz, 1H), 8.10 (s, 1H), 8.01 (dd, *J* = 8.1, 1.6 Hz, 1H), 7.91 (d, *J* = 8.0 Hz, 1H), 7.79 (d, *J* = 9.0 Hz, 1H), 7.74 (t, *J* = 8.0 Hz, 1H), 7.42 (t, *J* = 6.9 Hz, 1H), 5.16 (dd, *J* = 13.3, 5.2 Hz, 1H), 4.56 (d, *J* = 17.3 Hz, 1H), 4.44 (d, *J* = 17.4 Hz, 1H), 2.94 (ddd, *J* = 17.2, 13.7, 5.4 Hz, 1H), 2.68 – 2.59 (m, 1H), 2.48 – 2.39 (m, 1H), 2.05 (dtd, *J* = 12.7, 5.4, 2.3 Hz, 1H).

**Compound 9**

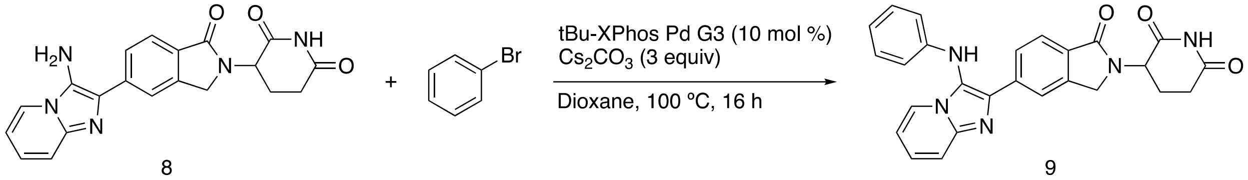


Compound 9 was synthesized following a reference procedure.[3] Compound 8 (9.0 mg, 24 µmol, 1.0 equiv) was mixed with cesium carbonate (23 mg, 72 µmol, 3.0 equiv) and bromobenzene (4.1 mg, 26 µmol, 1.1 equiv) in a glass vial. After three cycles of vacuum/nitrogen, the tBu-XPhos Pd G3 (1.9 mg, 2.4 µmol, 0.1 equiv) was added, and the reaction vessel was purged with vacuum/nitrogen again. Anhydrous dioxane (200 µL) was purged with nitrogen for at least 10 minutes and added to the reaction mixture. Nitrogen was bubbled through the mixture, and the vial was sealed and stirred at 100 ºC for 16 hours. Product formation was detected via LC-MS. The mixture was diluted in DMSO, filtered and purified via preparative HPLC using MeOH:$H_2O$ (0.035% formic acid additive) elution gradient. Yield: 5.1 mg, 47%. **LC-MS**: calculated exact mass = 451.16 found $[M+H]^+$ = 452.26. **$^1$H NMR** (500 MHz, DMSO-$d_6$) δ = 11.00 (s, 1H), 8.50 (s, 1H), 8.22 (s, 1H), 8.16 (d, *J* = 7.8 Hz, 2H), 7.82 (d, *J* = 7.8 Hz, 2H), 7.63 (m, 1H), 7.22 – 7.13 (m, 3H), 6.78 (t, *J* = 7.3 Hz, 1H), 6.62 (d, *J* = 7.9 Hz, 2H), 5.13 (dd, *J* = 13.3, 5.1 Hz, 1H), 4.51 (d, *J* = 17.3 Hz, 1H), 4.37 (d, *J* = 17.3 Hz, 1H), 2.92 (ddd, *J* = 17.3, 13.6, 5.4 Hz, 1H), 2.61 (m, 1H), 2.41 (m, 1H), 2.05 – 1.98 (m, 1H).

### NMR Spectra

#### $^1$H NMR of compound 1 (500 MHz, DMSO-d6)

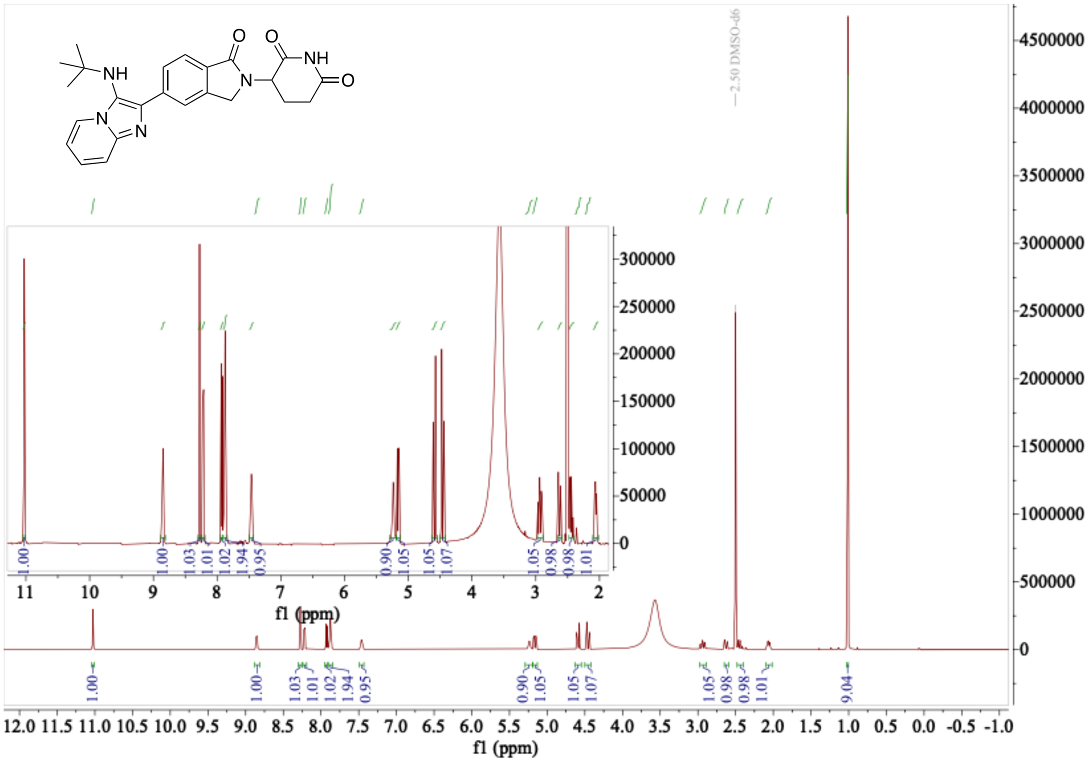

#### $^1$H NMR of compound 1-Me (500 MHz, DMSO-d6)

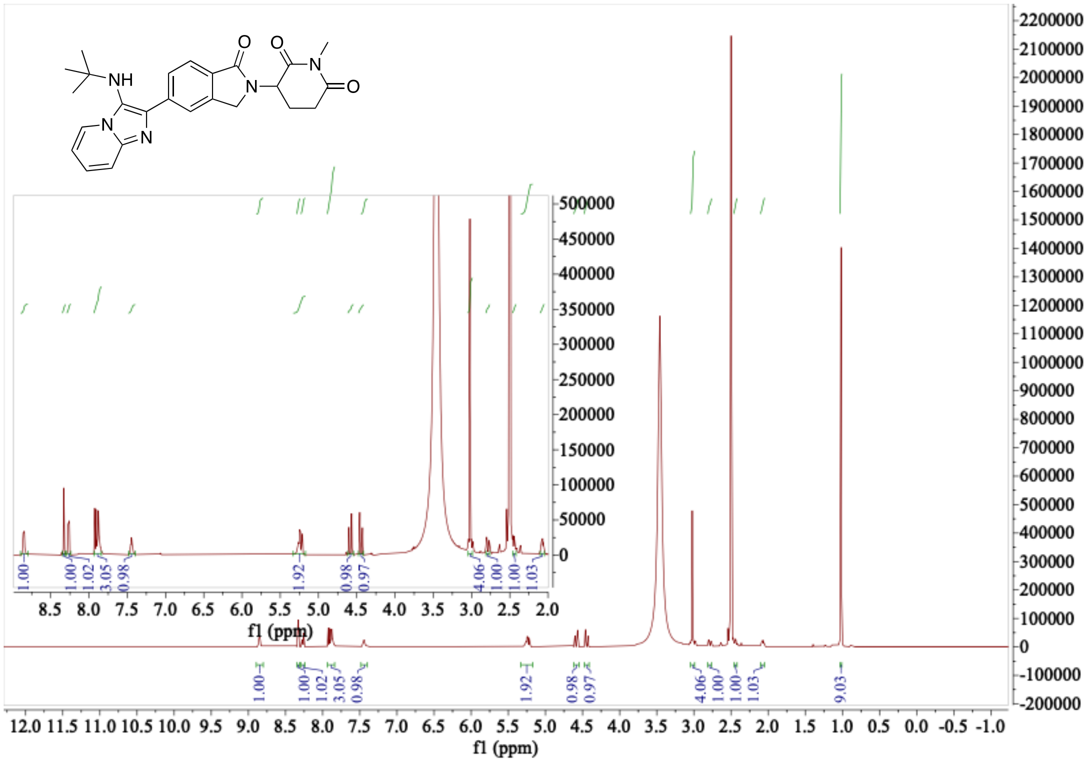

**[1]H NMR of compound 2 (500 MHz, DMSO-d6)**

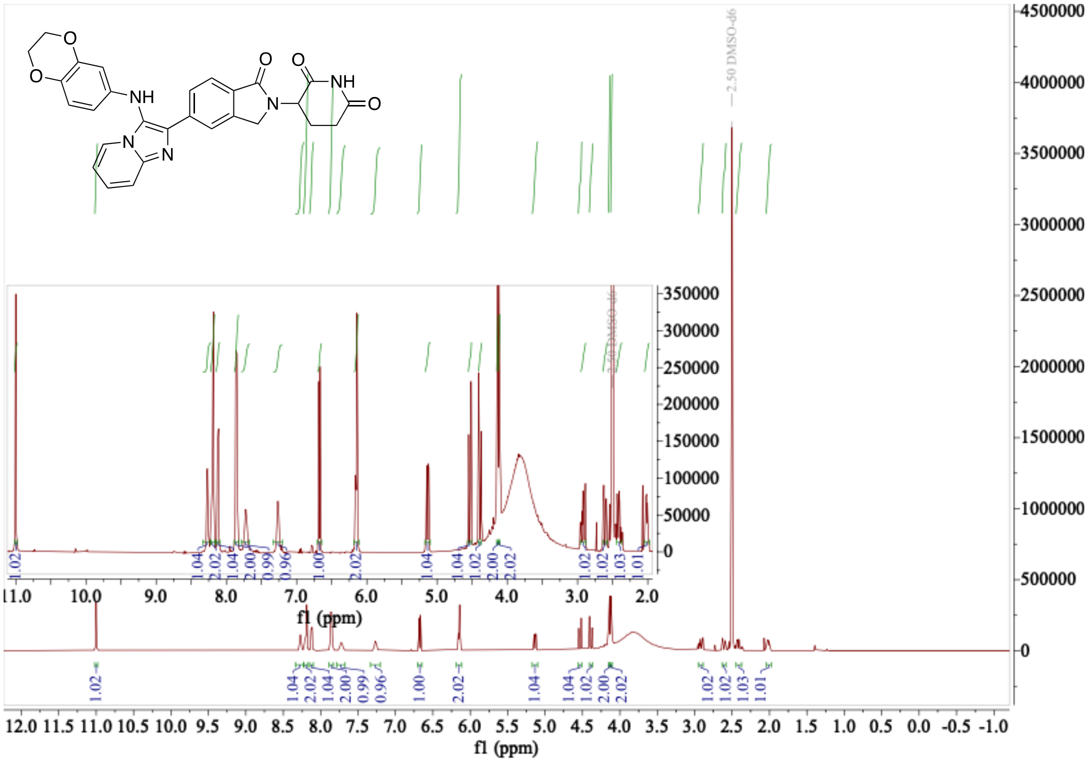

**[1]H NMR of compound 3 (500 MHz, DMSO-d6)**

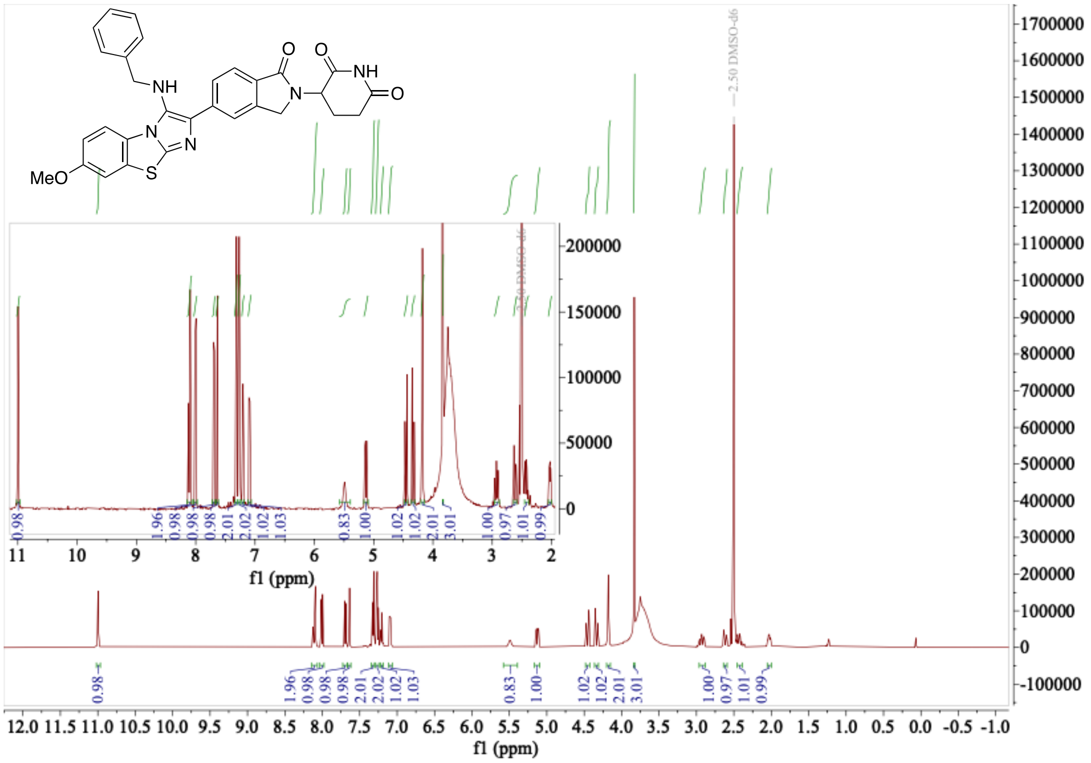

**[1]H NMR of compound 4 (500 MHz, DMSO-d6)**

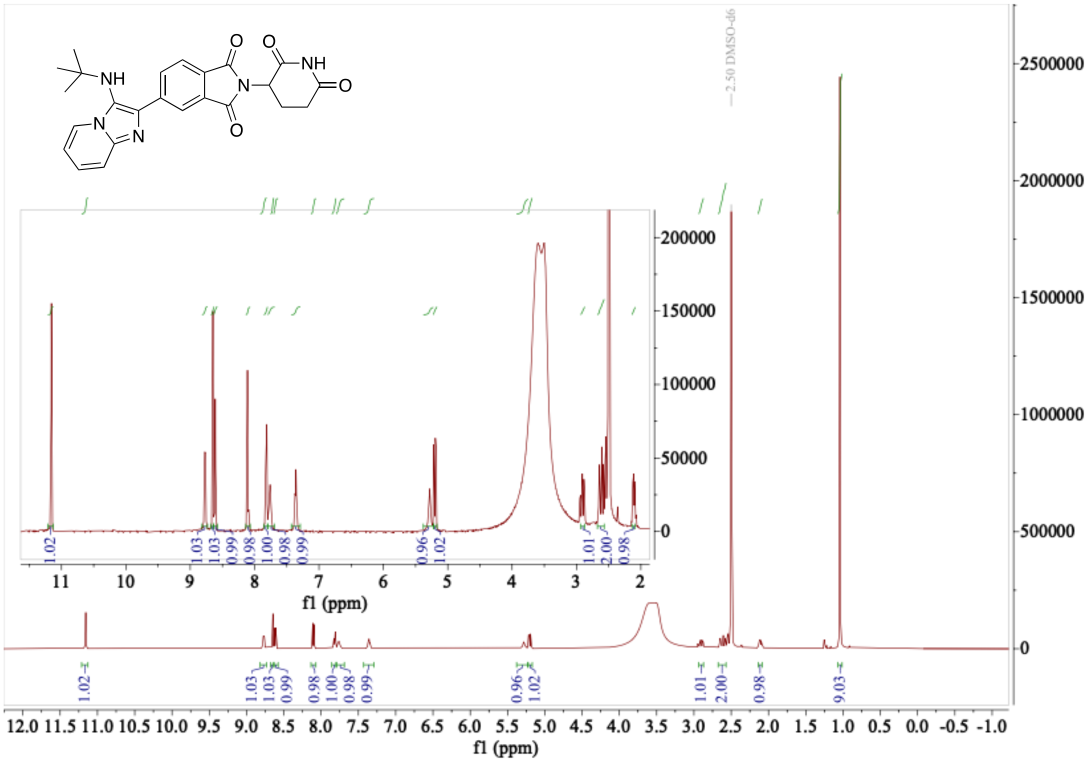

**[1]H NMR of compound 5 (500 MHz, DMSO-d6)**

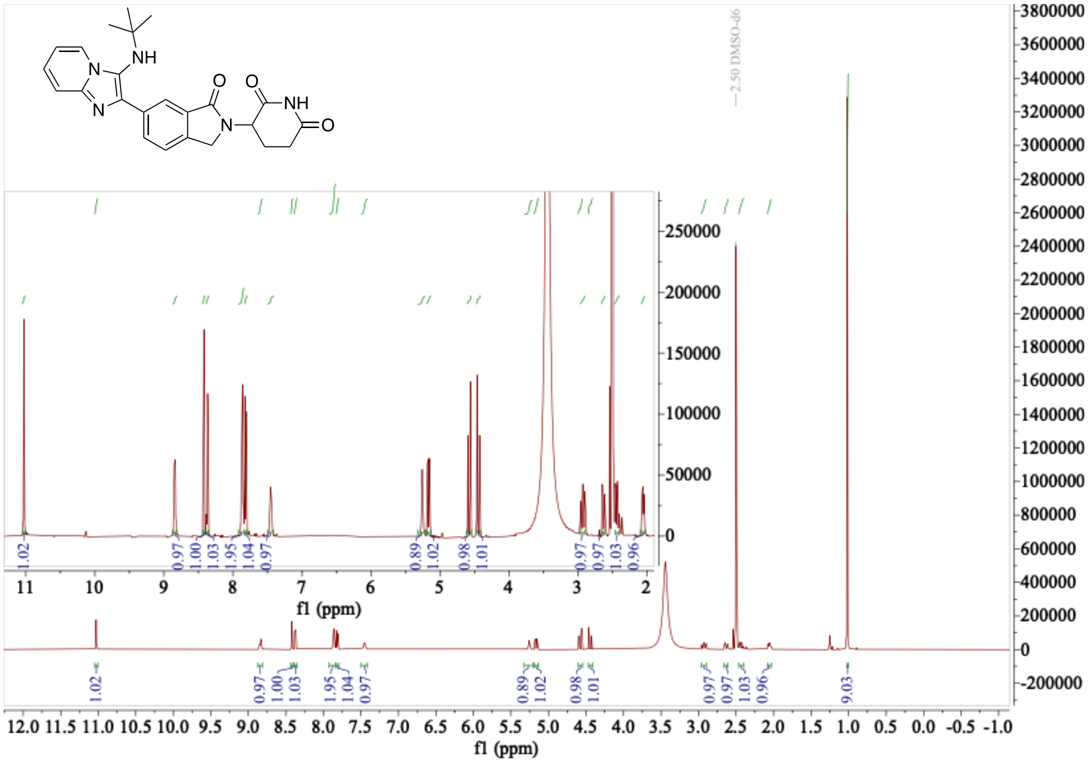

### $^1$H NMR of compound 6 (500 MHz, DMSO-d6)

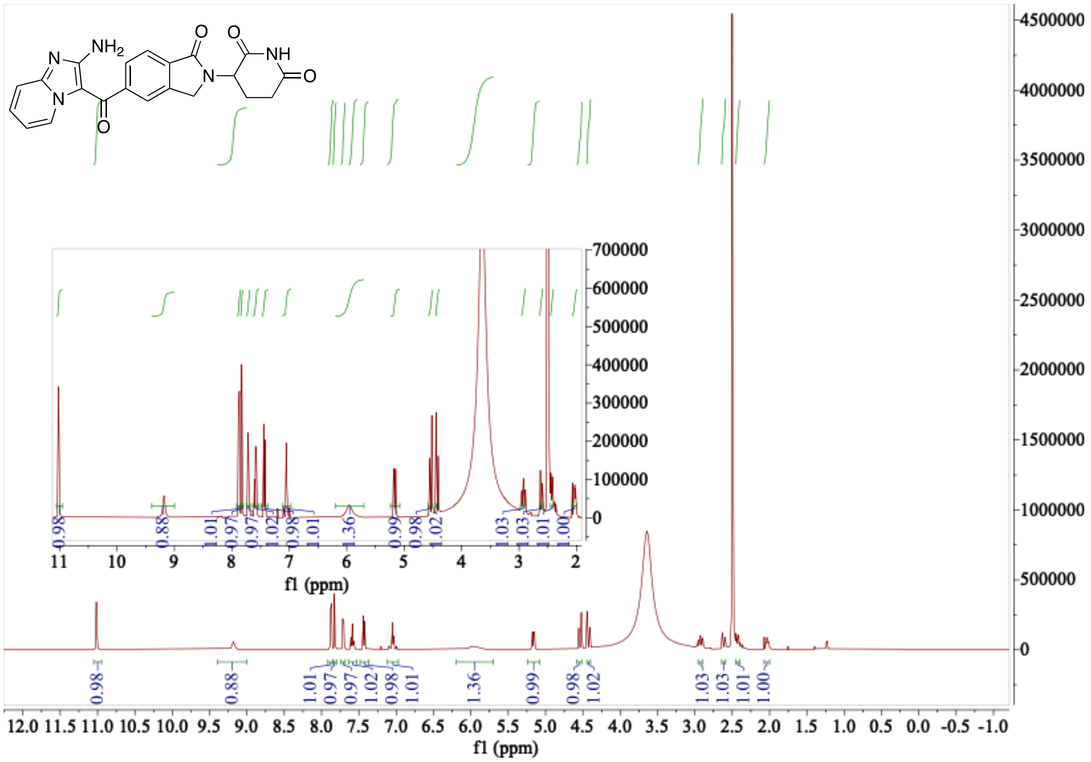

### $^1$H NMR of compound 7 (500 MHz, DMSO-d6)

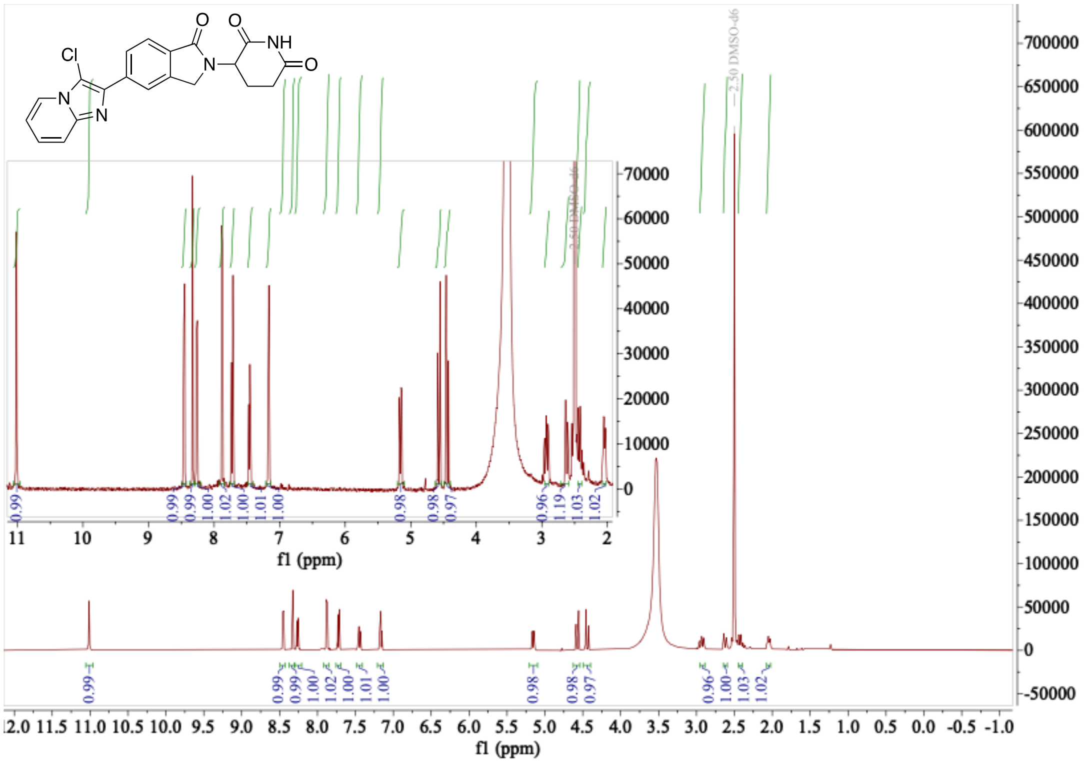

[1]H NMR of compound 8 (500 MHz, DMSO-d6)

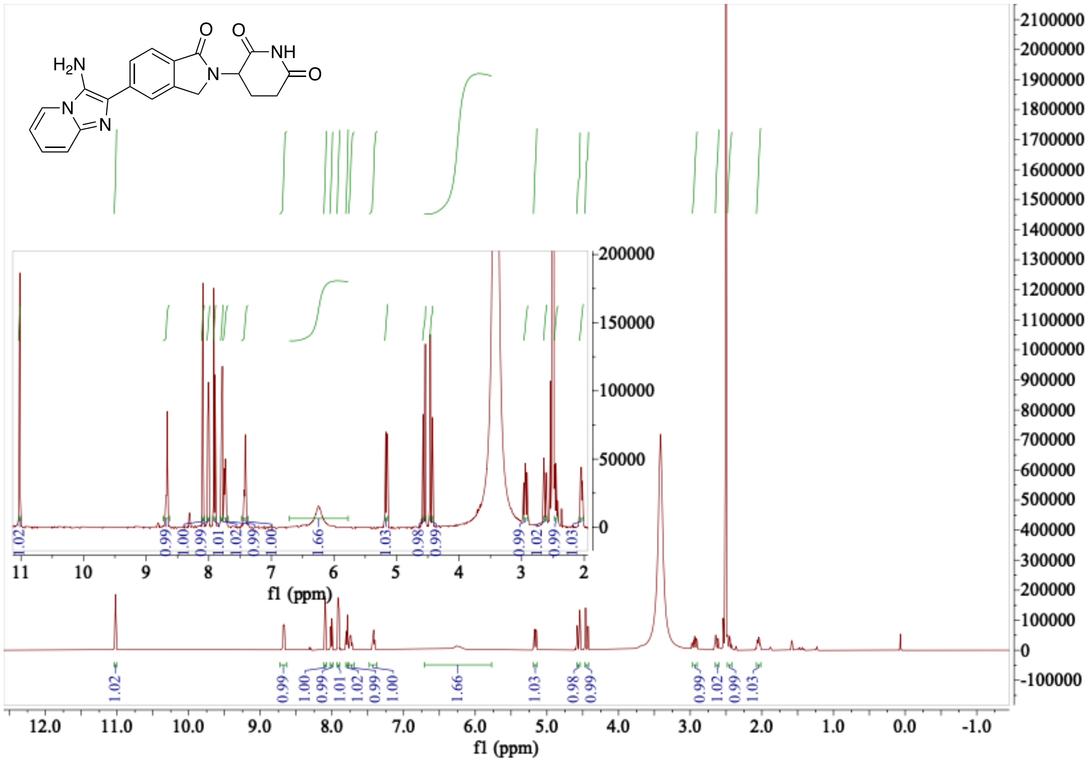

[1]H NMR of compound 9 (500 MHz, DMSO-d6)

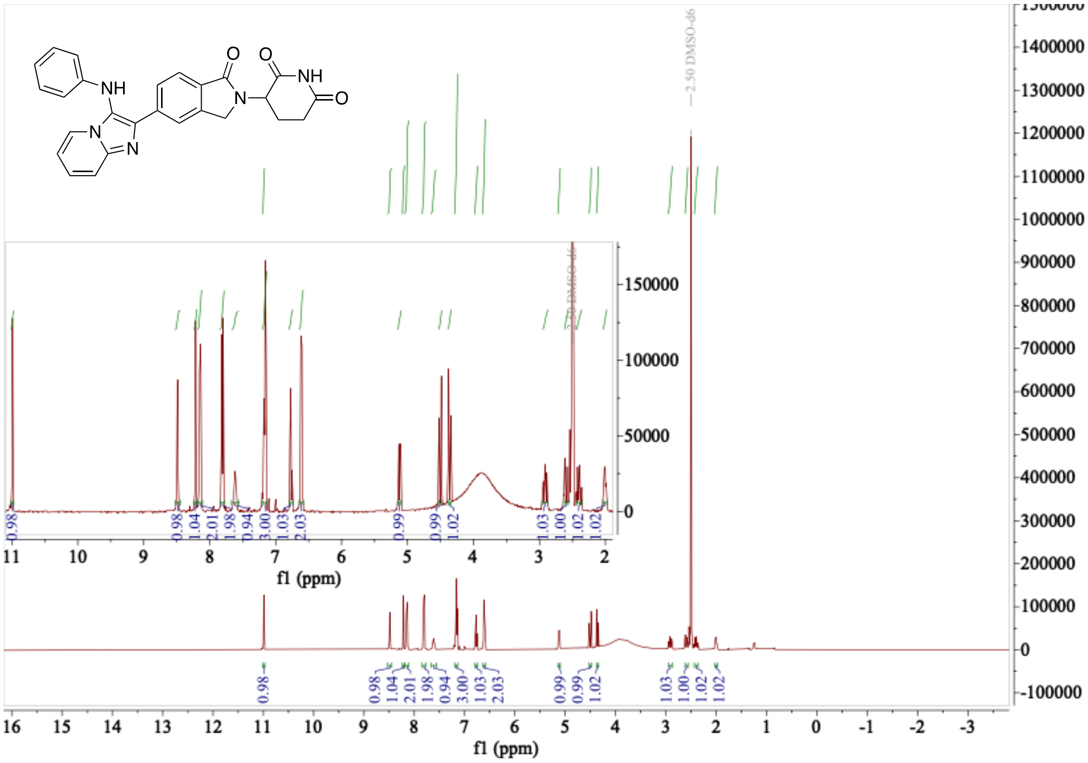

[1]H NMR of compound 10 (500 MHz, DMSO-d6)

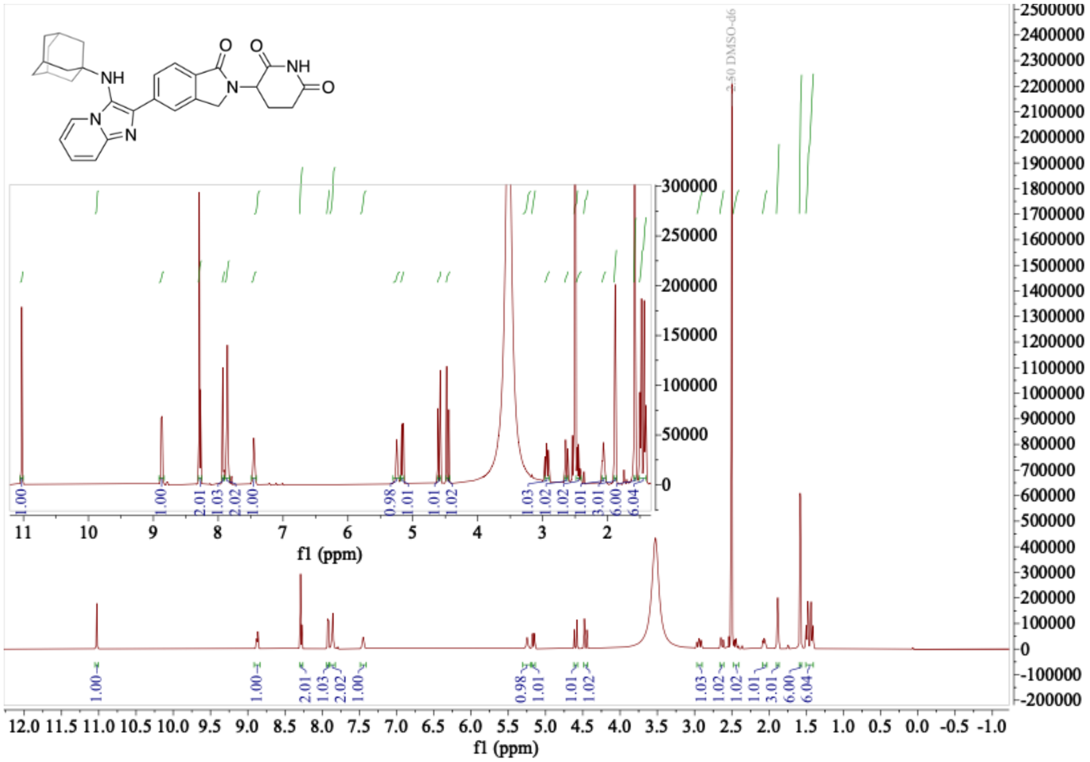

[1]H NMR of compound 11 (500 MHz, DMSO-d6)

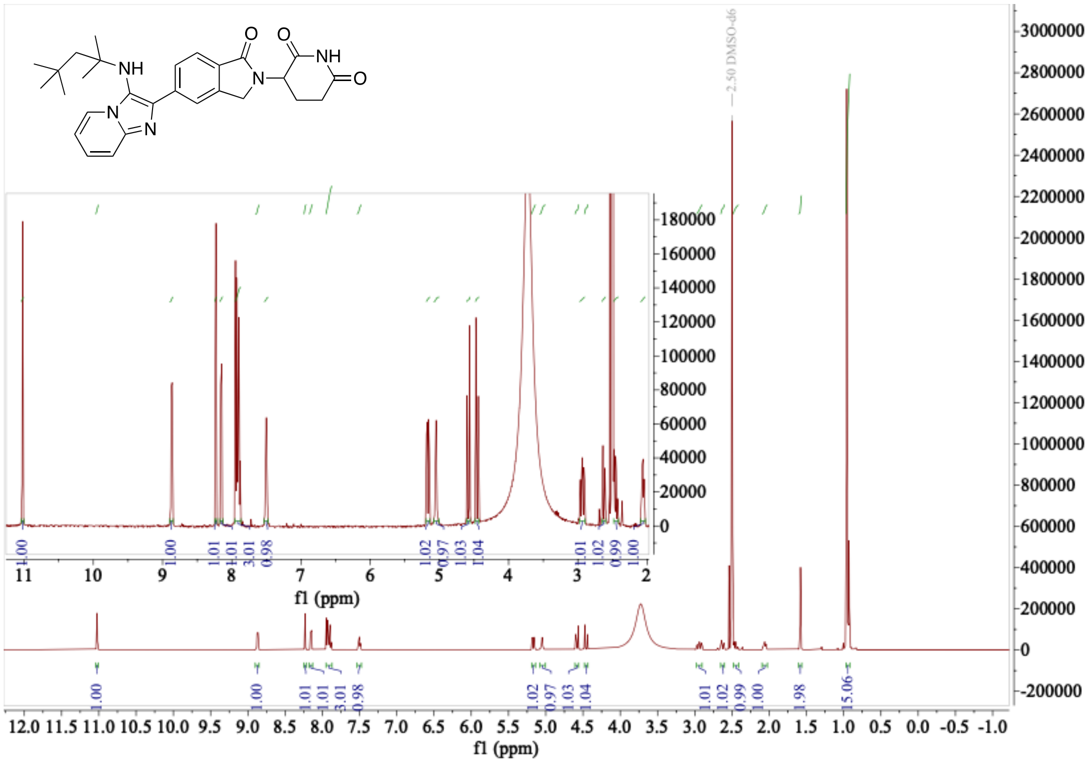

**¹H NMR of compound 12 (500 MHz, DMSO-d6)**

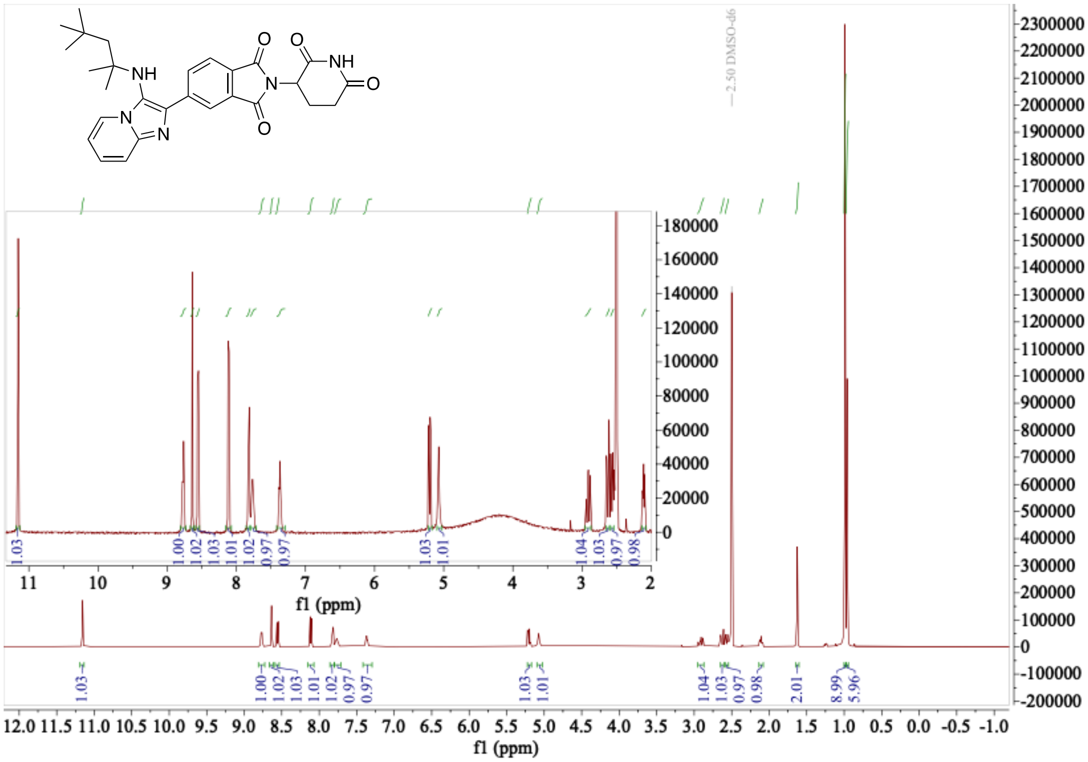

**¹H NMR of compound 13 (500 MHz, DMSO-d6)**

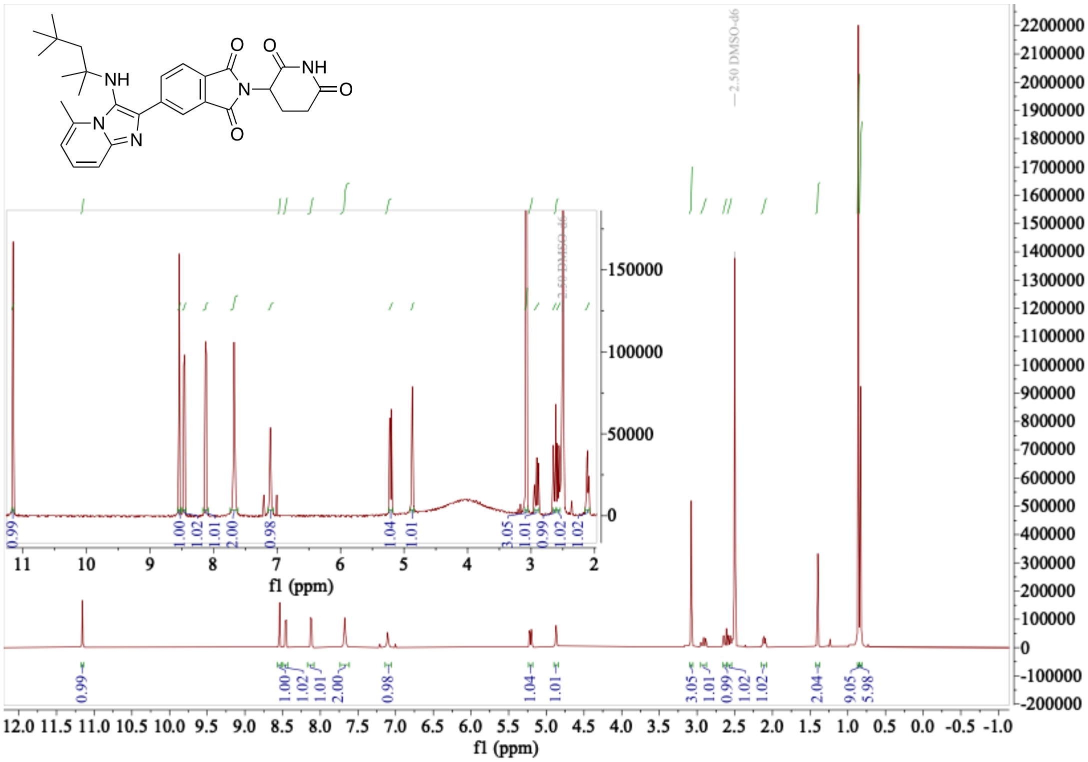
